# Morphological changes and two *Nodal* paralogs drive left-right asymmetry in the squamate veiled chameleon (*C. calyptratus*)

**DOI:** 10.1101/2023.01.18.524635

**Authors:** Natalia A. Shylo, Sarah E. Smith, Andrew Price, Fengli Guo, Melainia McClain, Paul Trainor

## Abstract

The ancestral mode of left-right (L-R) patterning involves cilia in the L-R organizer. However, the mechanisms regulating L-R patterning in non-avian reptiles remains an enigma, since most squamate embryos are undergoing organogenesis at oviposition. In contrast, veiled chameleon (*Chamaeleo calyptratus*) embryos are pre-gastrula at oviposition, making them an excellent organism for studying L-R patterning evolution. Here we show that veiled chameleon embryos lack motile cilia in their L-R organizer, consistent with the loss of motile cilia being a synapomorphy of all reptiles. Furthermore, in contrast to avians, geckos and turtles, which have one *Nodal* gene, veiled chameleon exhibits expression of two paralogs of *Nodal* in the left lateral plate mesoderm, albeit in non-identical patterns. Using live imaging, we observed asymmetric morphological changes that precede, and likely trigger, asymmetric expression of the Nodal cascade. Thus, veiled chameleons are a new and unique model for studying the evolution of L-R patterning.

## INTRODUCTION

Asymmetries are deeply rooted in the phylogenetic tree and are frequently driven by the ancient Nodal signaling cascade. For example, Nodal drives the asymmetric positioning of asexual buds in radially symmetric hydra ^1^. A duplication event resulted in two *Nodal* genes in gnathostomes, with additional expansion events in fish and frogs^2^. In contrast, mammals and avians retained a single copy of *Nodal* each, but different paralogs^2,3^. Henceforth we will refer to mammalian *Nodal* as *Nodal1*, and to avian *Nodal* as *Nodal2*, after Kajikawa et al. 2020^3^. The Nodal molecular cascade drives the establishment of left-right (L-R) asymmetry in deuterostomes, but what initiates asymmetric *Nodal* expression across phyla remains poorly understood. This is in part due to the constraints of the limited model organisms utilized.

A L-R organizer (LRO) is a transient organ in deuterostomes, and motile cilia and asymmetric cilia-generated flow have been shown to be a synapomorphy of deuterostome LROs, from sea urchins to humans^4^. Interestingly, avians, as well as some even-toed ungulates^5^ lack motile cilia in their LROs. Instead, asymmetric *Nodal* expression and L-R asymmetry are established through asymmetric cell movements and tilting of the LRO, which are tightly linked to primitive streak formation^5,6^. What triggers this asymmetric tissue morphogenesis currently remains unknown.

Non-avian reptiles comprise 40% of all living amniotes, yet their early development remains poorly understood. One of the major challenges is accessing early-stage reptilian embryos for detailed analysis. Access is often restricted to short breeding seasons, and further complicated by the fact that most squamate embryos are undergoing organogenesis at the time of oviposition, thus necessitating euthanasia of the mother for the study of L-R patterning events^7,8^. Thus, it was only recently revealed that turtles and geckos, like avian reptiles, lack motile cilia in their LRO^3^. However, unlike avians, no L-R asymmetric tissue morphogenesis was noted^3^. Furthermore, each retained only *Nodal2* in their genomes, suggesting that loss of motile cilia in the LRO and loss of *Nodal1* may be synapomorphies of all reptiles^3^.

Veiled chameleons (*Chamaeleo calyptratus*) breed year-round, depositing large clutches of eggs^9^. Most importantly, veiled chameleon embryos are pre-gastrula at oviposition, making them an excellent research organism for studying the mechanisms governing the regulation and evolution of gastrulation and L-R patterning ^10^. We discovered that in contrast to other reptiles described to-date, veiled chameleon embryos express both *Nodal1* and *Nodal2* in a L-R asymmetric manner. Furthermore, we found that veiled chameleons lack motile cilia in their LRO. Therefore, we used live imaging to track the timing and pattern of asymmetric tissue rearrangements to provide a deeper understanding of gastrulation and the establishment of L-R asymmetry in squamates. We find that asymmetric morphological changes occur prior to the onset of molecular L-R asymmetry. The tissue rearrangements that we observed differ from chicken and even-toed ungulate embryos, since non-avian reptiles lack a primitive streak, and instead use a blastopore and involution for gastrulation. Thus, veiled chameleons can serve as a new model for studying the evolution of gastrulation and L-R patterning.

## RESULTS

### Left-right patterning features in veiled chameleons

Embryonic turning is a distinct developmental event in the establishment of asymmetry during amniote embryogenesis. Previously it was assumed in chicken embryos, that asymmetric heart looping drove asymmetric embryo turning^11^. However, these processes can be de-coupled and are considered distinct independent developmental events ^12,13^. In veiled chameleon, embryo turning initiates with the head turning to the right at around the 4-somite stage (4ss) (Fig. 2 g, n, v), prior to the formation of the heart tube and subsequent heart looping. MF-20 staining reveals the formation of two heart fields at the 5-6ss (Fig. 1 a), which then fuse into a heart tube, that forms at an angle, pre-positioned for subsequent heart looping (Fig. 1 b). A heartbeat typically becomes detectable by the 10ss, concomitant with the heart undergoing looping morphogenesis (Fig. 1 c-e), resulting in a three-chambered organ (Fig. 1 f).

**Fig. 1.**
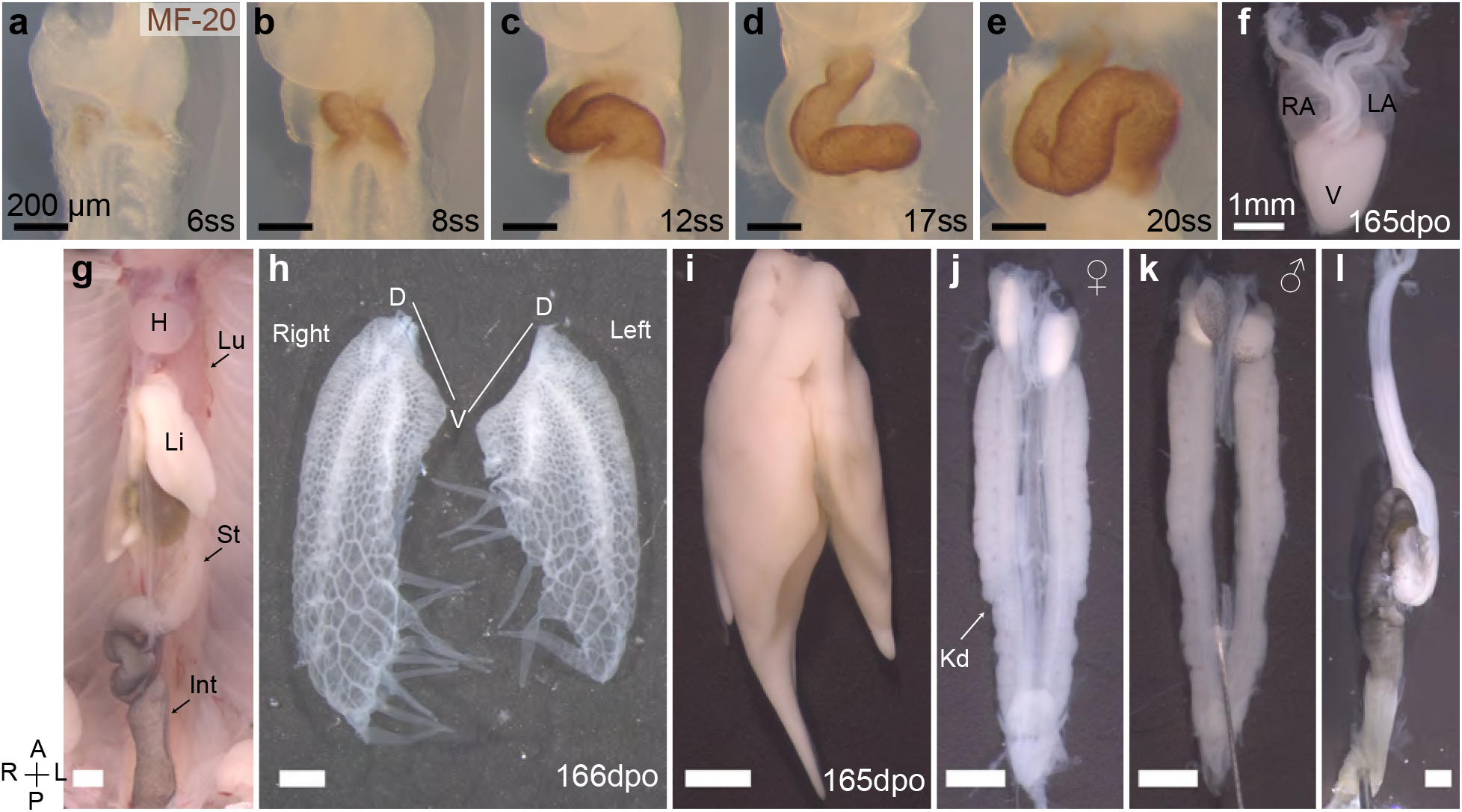
Organ asymmetry in veiled chameleon embryos. **a-f.** Progression of heart formation and development. In **a-e** the formation of the heart is highlighted through immunohistochemical staining for MF-20. **f.** The heart of pre-hatching stage embryos (165 dpo – days post oviposition) has three chambers – left and right atria (LA and RA, respectively) and one ventricle (V). **g.** General overview of abdominal organs in 165 dpo embryo. Heart (H), lungs (Lu), liver (Li), stomach (St), intestine (Int) are labeled. **h.** Lungs of a 166 dpo embryo have a single 3-chambered lobe on each side. The left lung is shorter than the right. Chameleons have unique diverticula on ventral ends of the lungs on both sides. Diverticula are arranged in two rows on the right lung, and in a single row on the left lung. The left lung has 5, and the right has 3+4 diverticula. **i.** The liver has a larger lobe on the right side. **j, k.** The urogenital system of both females (**j**) and males (**k**) exhibit L-R asymmetry with the left gonad being lower than the right. Kidneys (Kd) are elongated and lie flat against the dorsal abdominal wall. **l.** The gastrointestinal tract is relatively short, with stomach on the left side. The intestine posterior to the stomach has a single major loop and have black pigmentation. All panels show ventral view, unless otherwise noted. Anterior (A) is on the top, posterior (P) is on the bottom, left (L) is on the reader’s right, and right (R) is on the reader’s left. All black scale bars are 200 μm, all white scale bars are 1mm.

**Fig. 2.**
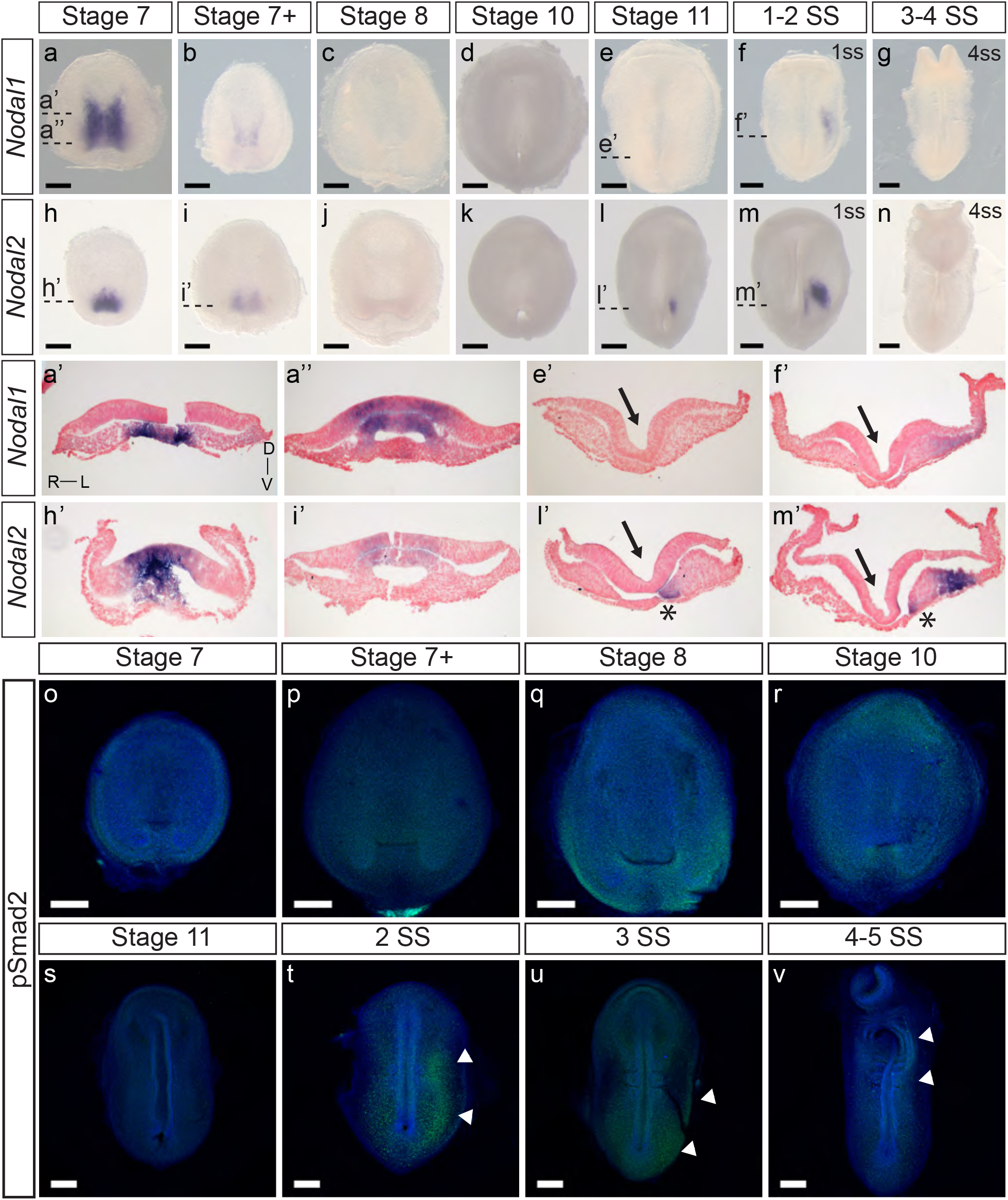
Expression and activity of *Nodal1* and *Nodal2* in veiled chameleon embryos. All embryos are presented in ventral view, unless otherwise noted. Dorsal view available in Figs. S2 and S3. **a-g.** Whole mount RNA *in situ* hybridization for *Nodal1* expression. **h-n.** Whole mount RNA *in situ* hybridization for *Nodal2* expression. **a’, a”, e’, f’, h’, i’, l’, m’**. Transverse sections of embryos from **a-n,** as indicated with dashed lines. Sections were counterstained with nuclear fast red for better visualization. Section orientation is as indicated in **a’**. Asterisk denotes *Nodal2* expression in presomitic mesoderm. Arrows point to the medial hinge point of the neural tube. **o-v.** The presence of active Nodal signaling was evaluated through antibody staining for pSmad2 (green), with nuclear Dapi staining in blue. White arrowheads denote areas of pSmad2 enrichment visible in this view. Normalized pSmad2 staining of embryos and intensity analysis between left and right sides is available in Fig. S3. All scale bars are 200 μm.

The heart is positioned high in the chest and is one of many structurally and positionally asymmetrical organs (Fig. 1 g). Veiled chameleon lungs have single 3-chambered lobes on each side (Fig. 1 h). Both lungs have structures of unknown function, unique to chameleons, called diverticula^14^. The left lung has a single row of five diverticula, whereas the right lung has two rows of three and four diverticula (Fig. 1 h). The left lung is notably shorter than the right lung. The liver has a larger lobe on the right side, similar to other animals (Fig. 1 i). The urogenital system is asymmetric in both sexes, with the axial position of the gonads and kidneys on the right being slightly more anterior than on the left (Fig 1 j, k). The gastrointestinal system is relatively short, with the stomach positioned on the left side, and a single characteristic loop of the intestine projecting towards the right side of the body (Fig. 1 g, l).

### *Nodal1* and *Nodal2* are both present in veiled chameleon transcriptome and are expressed asymmetrically during L-R patterning

Although a transcriptome of limited size was already available for veiled chameleons, it poorly correlated with the stages of development examined in this study^15^. Therefore, we collected a pool of embryos from St. 7 to 8 somites (Supplementary Dataset 1) and carried out Iso-seq (PacBio) to assemble a new transcriptome, which revealed the presence of most genes in the Nodal pathway and thus their expression during gastrulation and early somitogenesis stages. Notably, we identified two different transcripts for *Nodal*. Evolutionary comparison between predicted open reading frames and proteins from other animals revealed one of the transcripts as *Nodal*, and the other transcript as being *Nodal-like*. We adapted the nomenclature proposed by Kajikawa et al. 2020, and henceforth will refer to the former as *Nodal1* and the latter as *Nodal2* ^3^ (Fig. S1). Thus, contrary to a previous report that *Nodal1* was lost in reptiles^3^, we discovered that *Nodal1* was retained in veiled chameleons. Furthermore, subsequent genomic and phylogenetic analyses revealed several other squamate species have also retained *Nodal1* in their genomes (Fig. S1).

We next examined the expression pattern of *Nodal1* and *Nodal2* at the time of L-R establishment to infer their roles in this process (Fig. 2). Both *Nodal* transcripts exhibit a dynamic pattern of expression that suggests involvement in both gastrulation and L-R patterning. *Nodal1* and *Nodal2* are expressed symmetrically during gastrulation at stages 7-7+ in the blastopore (Fig. 2 a, b, h, i). In both cases, expression is in the chordamesoderm in the anterior region (Fig. 2 a’, h’) and in the chordamesoderm and blastopore plate more posteriorly (Fig. 2 a”, i’) ^16^. The differences in expression pattern are suggestive of distinct functions in gastrulation. With the onset of neurulation at stage 8 and appearance of the prechordal plate (Supplementary Dataset 1), the expression of both transcripts is extinguished, and remains absent through the early stages of the midline, head process and foregut formation (Fig. 2 c, d, j, k). *Nodal2* expression re-emerges just prior to the initiation of somitogenesis in a narrow domain just anterior to the blastopore slit in the left presomitic mesoderm (Fig. 2 l, l’). In a stage-matched embryo, *Nodal1* remains switched off (Fig. 2 e). With the onset of somitogenesis, *Nodal2* expression propagates to the left lateral plate mesoderm (LPM) in the posterior end of the embryo (Fig. 2 m, m’). Whether in response to *Nodal2*, or independent of *Nodal2*, *Nodal1* likewise becomes expressed in the LPM, in a much more diffuse manner (Fig. 2 f, f’). The expression of both *Nodal1* and *Nodal2* then expands in the left LPM but is downregulated and switched off by the 4ss (Fig. 2 g, n).

Smad2 becomes phosphorylated as the result of Nodal activity. For ease of analysis, we normalized pSmad2 expression against DAPI staining (Fig S3 i-p) and carried out an intensity analysis to compare left and right sides (Fig. S3 i’-p’). We do not observe specific enrichment of pSmad2 near the blastopore during the late stages of gastrulation, suggesting that Nodal1/2 molecules may act through a different pathway at this stage (Fig. 2 o-s, S3 i-m, i’-m’). With the onset of somitogenesis, broad pSmad2 expression becomes apparent in the left LPM, and lingers until the 4ss, slightly later than the expression of *Nodal1/2* (Fig. 2 t-v, S3 n-p, n’-p’). Notably, we did not observe an enrichment of pSmad2 in the presomitic mesoderm, in contrast to a previous report about the Madagascar ground gecko^3^.

### The expression patterns of key members of the Nodal cascade align with *Nodal1/2* expression

Characterizing the dynamic expression of *Nodal1* and *Nodal2* and their activity allowed us to narrow down the timing of L-R patterning events in veiled chameleons. We then set out to identify and examine the expression patterns of other genes in the Nodal cascade. We identified a single *Lefty* gene in veiled chameleons as is typical for most vertebrates^17^. Lefty is a well-known repressor of Nodal. In chicken and mouse embryos, *Lefty* is expressed in the LPM and the midline, acting both as a midline barrier, and repressor of *Nodal* in the LPM through an activator-inhibitor loop^18^. Prior studies have suggested that midline *Lefty* expression is more critical for *Nodal* inhibition than its LPM expression, and under certain conditions *Lefty* expression may be downregulated without disrupting the signaling cascade^18,19^. *Lefty* expression in veiled chameleon embryos provides *in vivo* evidence for this model. In veiled chameleons *Lefty* is expressed exclusively in the midline (Fig. 3 a-g). It has strong symmetrical expression in the notochordal plate at stage 8, as *Nodal* becomes downregulated (Fig. 3 c, c’). The symmetrical midline expression of *Lefty* then becomes extinguished (Fig. 3 d, d’), followed shortly by its reexpression exclusively in the left side of the notochordal plate, which is distinct from its expression in left side of the floor plate of mouse and chicken embryos ^20,21^ (Fig. 3 e, e’). *Lefty* is initially expressed just anterior to the blastopore, at the axial level of early *Nodal2* expression, and then propagates anteriorly through the entire length of the notochordal plate, before being switched off by 4-5ss (Fig. 2 l, Fig. 3 e, e’, f, f’, g and not shown). The lack of *Lefty* expression in the LPM, therefore leaves the system without an off switch for *Nodal1/2* expression in the LPM.

**Fig. 3.**
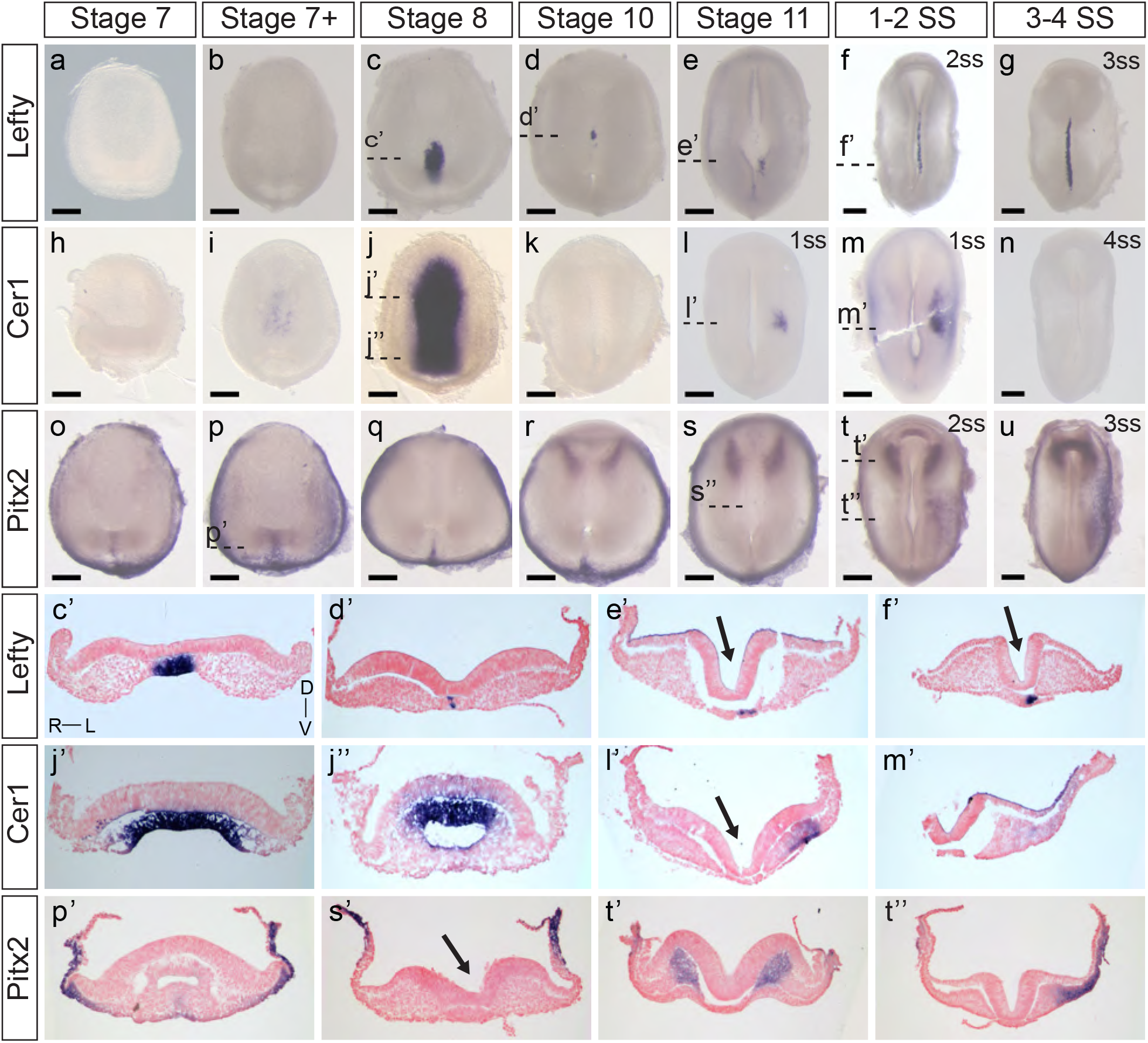
Expression patters of key members of the Nodal cascade in veiled chameleon embryos. All embryos are presented in ventral view, unless otherwise noted. Dorsal view available in Fig. S4. **a-g.** Whole mount RNA *in situ* hybridization for *Lefty* expression. **h-n.** Whole mount RNA *in situ* hybridization for *Cer1* expression. **o-u.** Whole mount RNA *in situ* hybridization for *Pitx2* expression. **c’, d’, e’, f’, j’, j”, i’, m’, p’, s’, t’, t”**. Transverse sections of embryos from **a-u,** as indicated with dashed lines. Sections were counterstained with nuclear fast red for better visualization. Section orientation is as indicated in **c’**. Arrows point to the medial hinge point of the neural tube. All scale bars are 200 μm.

*Dand5(Cerl2/Coco/Charon)* is the first asymmetrically expressed member of the Nodal cascade and acts to regulate the repression of Nodal^22^. Interestingly, we found no evidence for *Dand5* expression in the veiled chameleon transcriptome, and it appears to have been lost in all reptilian clades^3^. Since *Cer1* acts as the asymmetric repressor of *Nodal2* in chicken embryos, and belongs to the same family as *Dand5*, we focused on the pattern of *Cer1* expression instead^21,23^. *Cer1* is not expressed in veiled chameleon embryos prior to stage 8, and the lack of expression in the head region suggests it might play a lesser role in the anterior visceral endoderm than in other species^23^. However, we observed a huge, but temporary burst of *Cer1* in the chordamesoderm at stage 8 that coincides with downregulation of both *Nodal* genes (Fig 2 c, j, Fig. 3 j, j’, j”, k). *Cer1* is re-expressed later in the LPM, in a pattern reminiscent of *Nodal1* expression (Fig 2 f, f’, Fig. 3 l, l’, m, m’). *Cer1* expression is then downregulated by the 4-somite stage, similar to *Nodal1/2*. Overall, the dynamic pattern of *Cer1* expression is consistent with it being induced by *Nodal*, and then acting as a strong repressor of *Nodal1/2*, similar to its activity in chicken embryos^24^.

*Pitx2* serves as a critical transcriptional regulator of L-R asymmetry in vertebrates^25–27^ and we observed that *Pitx2* is expressed broadly in gastrulating veiled chameleon embryos (Fig. 3 o-r, p’). Subsequently, *Pitx2* expression becomes mostly restricted to the head mesoderm, as in other animals ^25^ (Fig. 3 r, s, t, t’, u), and the left LPM (Fig. 3 t, t”, u). Thus, the core of the Nodal signaling cascade is conserved in veiled chameleons, and the observed changes provide a framework to understand the mechanics of the process, as well as the opportunity for comparative analyses of gene and protein function evolution.

### Veiled chameleon lacks motile cilia at the left-right organizer

Most deuterostomes examined to-date, ranging from sea urchins to humans, use motile cilia in the LRO to establish L-R asymmetry^28,29^. Transcription factor *Foxj1* induces the formation of motile cilia, and its loss leads to L-R patterning defects^30^. *Dnah11* is an axonemal dynein, required for ciliary motility, and is mutated in the classical L-R patterning mouse mutant *inversus viscerum* (*iv*)^31^. Interestingly, we could not detect expression of *Foxj1* and *Dnah11* in veiled chameleons during the establishment of L-R patterning (Fig. S5 a-e, g-k). Both *Foxj1* and *Dnah11* are however expressed at later stages of veiled chameleon development, most notably in the otic vesicle (arrowheads in Fig S5 f and l), and kidneys (arrows in Fig. S5 f and l). The kidneys of many animals are known to contain motile cilia^32^, and we confirmed the presence of motile cilia in veiled chameleon kidneys through direct imaging of dissociated cells from the kidneys (Fig. S5 v, Supplementary video 1). We also verified our findings by performing SEM and TEM analyses of Stage 7+, 10 and 2ss veiled chameleon embryos especially near the blastopore, the presumed LRO^3^. We found no evidence of motile-like cilia via SEM but observed short primary cilia throughout the embryo (Fig. S5 m-r). TEM analysis of several cilia from the blastopore of stage 7+ embryos (Fig. S5 s) revealed all the cilia had a non-motile 9+0 conformation with no evidence for axonemal dyneins (Fig. S5 s-u), thus confirming the absence of motile cilia in the LRO of veiled chameleons.

### Morphological changes in the embryo temporally correlate with establishment of L-R asymmetry

Similar to veiled chameleons, avians and even-toed ungulates also lack motile cilia in their LROs^5^. Instead, L-R asymmetry is established in these species when cells surrounding the node migrate in an asymmetric manner, such that the node becomes tilted^5,6,33^. In avians, the asymmetric morphology of the node is well established^33^, but the asymmetric cell migration was revealed more recently through live imaging ^5,6^. Therefore, we evaluated chameleon embryonic development via live imaging to detect differences between the left and right sides in the establishment of L-R asymmetry. Veiled chameleons do not have a primitive streak or a distinct node, like avians or mammals, but have a blastopore slit instead^34^. However, despite the morphological differences between chickens and chameleons, their patterns of gastrulation appear strikingly similar, with polonaise movements clearly visible in gastrulating chameleon embryos (Supplementary video 2).

We successfully visualized the progression of L-R patterning through multiple timelapses of overlapping stages of development (Fig. 4, Supplementary videos 2-4). Transverse optical sections (O.S.) revealed dynamic morphological changes in the embryos, which were consistent between different samples (Fig. 4, Supplementary videos 2-4). As veiled chameleon embryos advance to stage 10 of development, we observed a subtle dorsal displacement of the left side of the embryo (Fig. 4 c’, d’). This morphological change becomes very pronounced by stage 11, when the left side has shifted much farther dorsally than the right side (Fig. 4 e’, g’ arrows). Subsequently, by the onset of somitogenesis, the right side of the veiled chameleon embryo moves dorsally to match the left side (Fig. 4 f’, h’ arrows). After that, both sides of the embryo appear to develop similarly from a dorsoventral morphological perspective. Notably, the earliest detectable morphological changes at stage 10 (Fig. 4 c, d, c’, d’) occur prior to the asymmetric *Nodal* expression (Fig. 2 d, k).

**Fig. 4.**
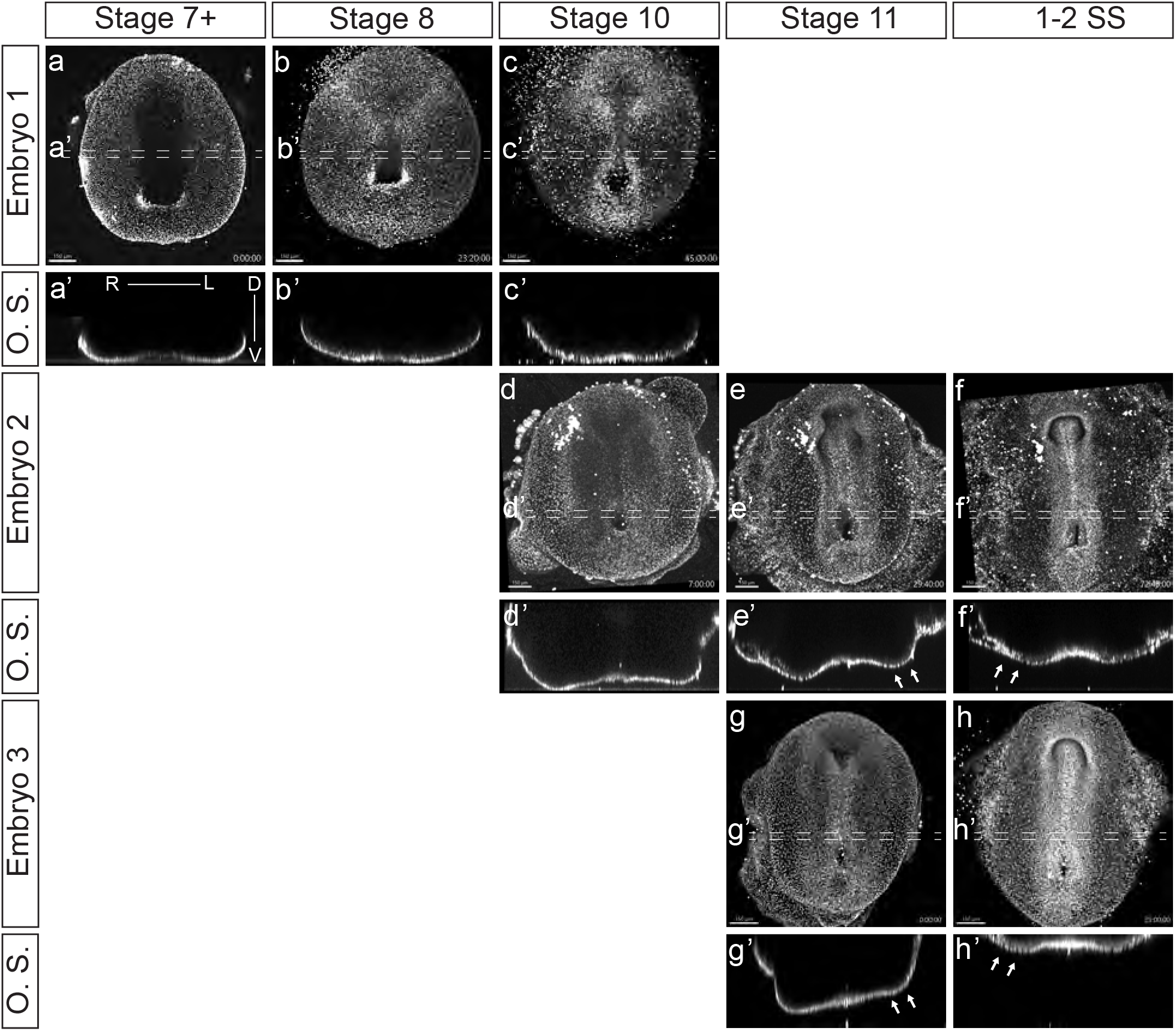
Video visualization of cellular migrations during gastrulation and left-right patterning. **a-h** Still images of developing embryos from videos 2-4 show ventral view of embryos cultured with 5-SiR-Hoechst STED DNA dye to label nuclei. The punctate lines show the approximate region of the 40 μm optical section. **a’-h’** Transverse optical sections (O.S) of embryos in **a-h**, from the regions, indicated by punctate lines. Section orientation is as indicated in **a’**. Videos 1-3 spanned different developmental stages, approximately assigned in the figure. White arrows in **e’, f’, g’, h’** show the areas of significant morphological changes in a developing embryo. Scale bars are in the bottom left corner of images and are 150 μm.

To circumvent the limitations of recording the development of embryos in culture with vital dyes, and to gain a better understanding and resolution of the processes that are taking place, we examined embryonic morphology in greater detail with RNA *in situ* markers and histological stains.

### Breaking L-R symmetry results from morphological changes during embryogenesis

To augment and further validate our live *ex ovo* imaging observations of the establishment of L-R asymmetry in veiled chameleon embryos, we performed expression analyses with classic genetic markers of morphological asymmetry in amniotes. We initially evaluated the expression of *Shh* (Fig. 5 a-g, o, d’-f’), since it is recognized as one of the first genes to be expressed asymmetrically in chicken embryos^35^. *Shh* expression first appears around stage 7+ in the prechordal plate of veiled chameleon embryos (Fig. 5 b-d, d’). Then as the midline develops, *Shh* expression becomes prominent in the floor plate and the notochordal plate (Fig. 5 d-g, d’, e’, f’) similar to other vertebrates. Notably, *Shh* expression in the notochord and floor plate at all times and at all levels in the embryo is L-R symmetrical (Fig. 5 d’, e’, f’).

**Fig. 5.**
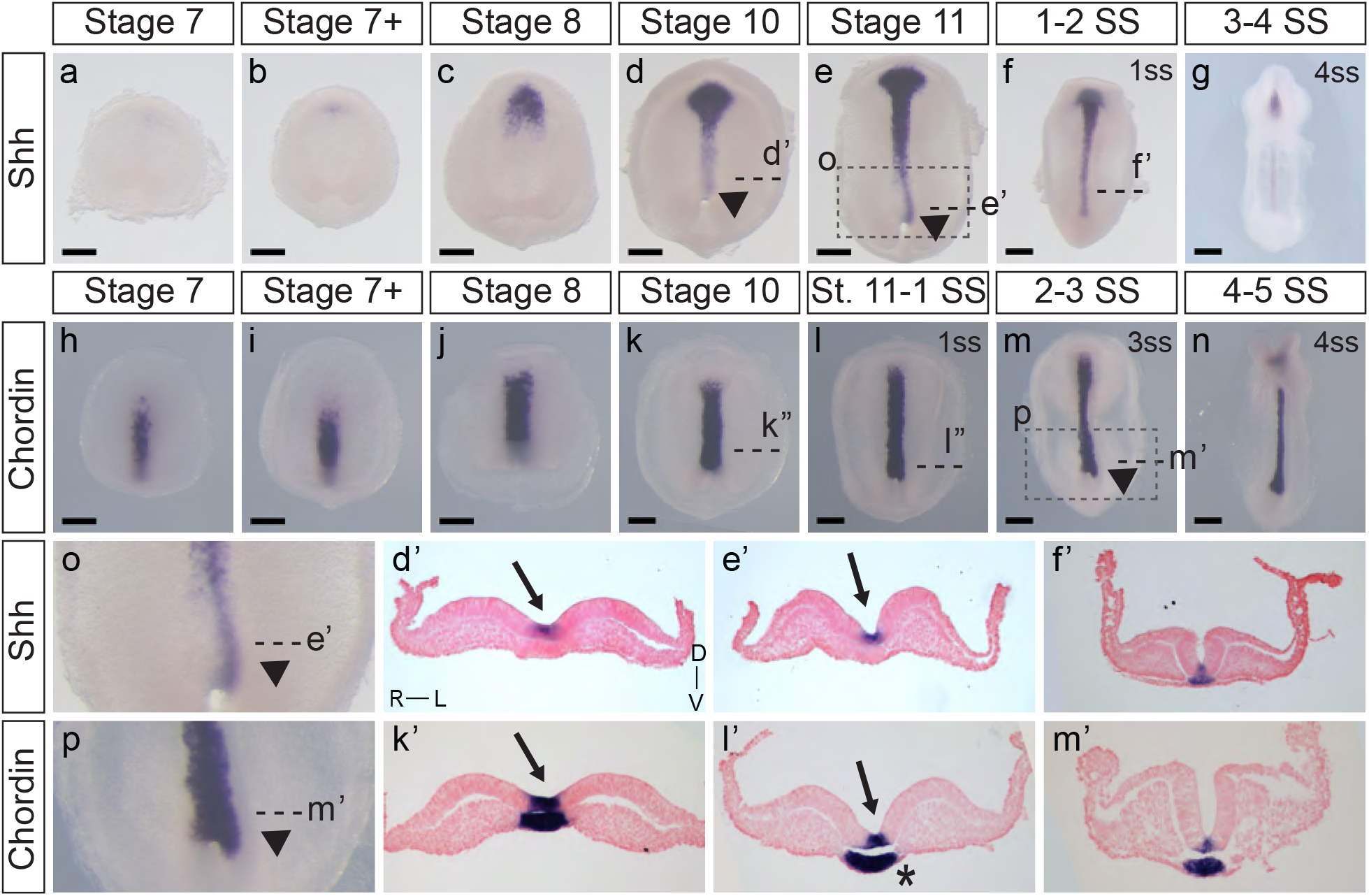
Expression of *Shh* and *Chordin* in veiled chameleons reveal tissue rearrangements and morphological changes in embryos. All embryos are presented in ventral view, unless otherwise noted. Dorsal view available in Fig. S7. **a-g.** Whole mount RNA *in situ* hybridization for *Shh* expression. Arrowheads point to asymmetric patterning of the midline near the blastopore. **h-n.** Whole mount RNA *in situ* hybridization for *Chordin* expression. **o, p.** Enlarged areas of asymmetry, as indicated in **e** and **m** with a dashed box. **d’, e’, f’, k’, l’, m’**. Transverse sections of embryos from **a-n,** as indicated with dashed lines. Sections were counterstained with nuclear fast red for better visualization. Section orientation is as indicated in **d’**. Arrows point to the medial hinge point of the neural tube. Asterisk reveals asymmetry in notochordal plate. All scale bars are 200 μm.

In chicken embryos, *Fgf8* is expressed asymmetrically in the posterior perinodal region complementary to *Shh* and is believed to maintain asymmetric gene expression and repression on the right side of the embryo^36,37^. Likewise, *Fgf8* is expressed in the posterior of veiled chameleon embryos in a pattern complementary to *Shh* (Fig. S7 o-bb). The onset of expression in chameleon is temporally similar to *Shh*, around stages 7-7+, becoming more pronounced in the posterior of the embryo throughout development (Fig. S7 o-bb, r’, s’, t’, u’). Similar to *Shh* expression, *Fgf8* is expressed symmetrically near the blastopore (Fig. S7 t’, u’).

When examining histological sections, we noted that the notochordal plate may also have L-R asymmetry in chameleon embryos (Fig. S7 s’ asterisk). We therefore used *Chordin* as an additional marker that labels both the floor plate and notochordal plate in veiled chameleon embryos in order to better examine notochordal plate morphology (Fig. 5 h-n, k’-m’). Its expression is strong at all timepoints examined and appears L-R symmetrical in both the floor plate and notochordal plate (Fig. 5 k’-m’).

Although *Shh, Fgf8* and *Chordin* are each expressed L-R symmetrically, as evident in transverse sections (Fig. 5, Fig. S7), whole mount *in situ* hybridization staining revealed remarkable morphological L-R asymmetry of *Shh* and *Chordin* (Fig. 5), and, to a lesser degree *Fgf8* (Fig. S7), in the posterior region of the embryos, phenotypes most striking for the midline markers (Fig. 5 d, e, o, l, m, p arrowheads, Fig. S7 d, e, l, m, t, aa). The asymmetry is most pronounced near the blastopore, with a deviation to the left (Fig. 5 d, e, o, l, m, p arrowheads, Fig. S7 d, e, l, m, t, aa). This morphological asymmetry is transient and disappears with the onset of somitogenesis. It is important to emphasize that *Shh, Chordin* and *Fgf8* are not themselves expressed asymmetrically, but serve here as midline markers that highlight the morphological asymmetry (Fig. 5, Fig. S7).

Transverse sections of stage 10, 11 and early somitogenesis embryos revealed the underlying morphological changes. The median hinge point of the forming neural tube is tilted to the left, instead of being centrally located (Arrows in Fig. 2, 3, 5, S7). The notochordal plate likewise displays L-R morphological changes (Asterisk Fig. 5 l’, S7 s’). Lastly, transverse sections of stage 10-11 and early somite embryos reveal the presence of more mesoderm on the left side of the embryo, compared to the right side (Most prominent in Fig. 2 f’, l’, m’ and Fig. 3 t”). To confirm that these observations were not an artifact of dissected and processed embryos, we performed paraffin sectioning of additional embryos at stages 10 and 11 with all their embryonic and extraembryonic tissues intact (Fig. S7 cc, dd). Hematoxylin and Eosin staining of transverse sections clearly reveal the displaced medial hinge point, and its asymmetric positioning in relation to the notochordal plate (Fig. S7 cc’, dd’).

Our observations from time lapse imaging and mRNA expression analyses corroboratively reveal that the onset of morphological changes in the embryo that lead to the establishment of L-R asymmetry, first occurs at stage 10, at a time when neither of the *Nodal* genes is expressed. Therefore, we conclude that morphological changes in the embryo are central to the mechanism that triggers molecular L-R asymmetry in veiled chameleon embryos.

## DISCUSSON

In this study we carried out a detailed developmental analysis of the breaking of L-R symmetry in veiled chameleon embryos and the mechanical and molecular mechanisms underpinning the process. Non-avian reptiles do not have a conventional node, and instead use a blastopore slit for gastrulation, which likely preforms the functions of the LRO^3,38^. Although motile cilia are central to the breaking of L-R symmetry in many vertebrate species, avians, turtles and geckos lack motile cilia in their LROs^3^. Similarly veiled chameleons also lack motile cilia in their LRO as L-R patterning becomes established. Thus, our study lends further support to the idea that the absence of motile cilia in the LRO is a synapomorphy of all reptiles.

While it was previously reported that *Nodal2* is the only *Nodal* ortholog conserved in reptiles^3^, we discovered that veiled chameleons possess two *Nodal* genes. We also found through genomic and phylogenetic comparisons that several other squamates also retained both *Nodal* genes in their genomes. In veiled chameleon, the expression of both orthologs is highly dynamic. *Nodal1* and *Nodal2* both appear to be involved in gastrulation and in L-R patterning but have distinct patterns of expression during gastrulation in different domains near the blastopore. Remarkably, the expression of both *Nodal* genes disappears abruptly at stage 8, coinciding with a strong burst of *Cer1* expression, symmetrical in the midline, together with *Lefty* (Fig. 6). The timing of these events implies that similar to observations in other species, Cer1 and Lefty are indeed inhibitors of Nodal signaling in veiled chameleons. Furthermore, their re-activation subsequent to *Nodal2* expression in the left LPM is consistent with an activator-inhibitor model of gene regulation^18^.

**Fig. 6.**
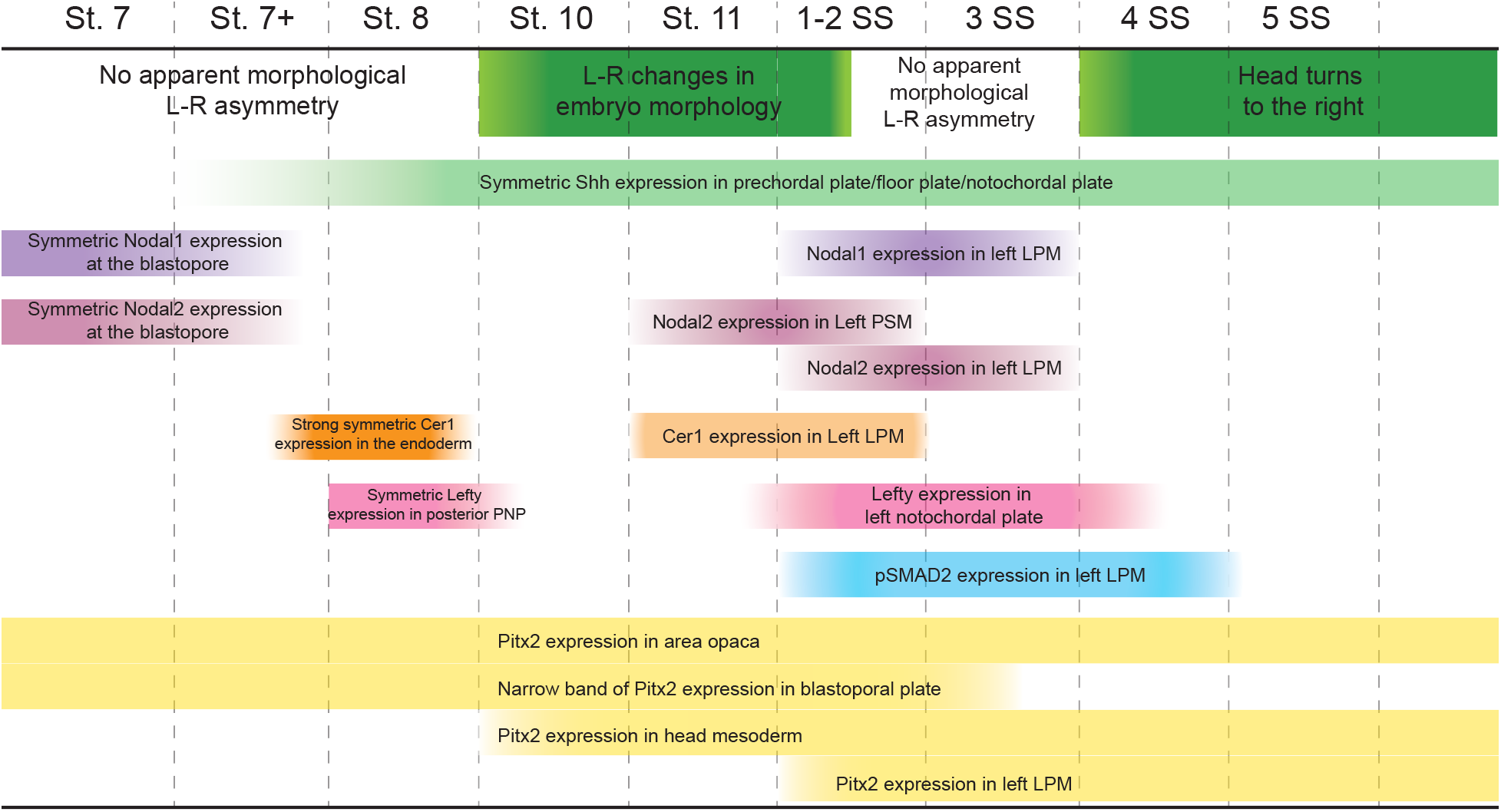
Timing of Nodal cascade markers expression and morphological changes in veiled chameleons. Temporal changes in the embryo set in the context of changing molecular markers expression, their pattern and intensity in the Nodal cascade. Left-right (L-R), lateral plate mesoderm (LPM), presomitic mesoderm (PSM), prospective notochordal plate (PNP).

Typically, Lefty acts as a Nodal inhibitor in the left LPM. However, veiled chameleons lack *Lefty* expression in the left LPM. Thus, we speculate that the veiled chameleon *Lefty* gene may have been subject to significant regulatory changes. It is interesting to note however, that under circumstances of low *Nodal* expression, the presence of Lefty in the LPM may be dispensable in the establishment of L-R asymmetry^19^. *Cer1* is an alternative Nodal repressor. It belongs to the same family of genes as *Dand5*, a known *Nodal* repressor that has been lost from reptile genomes^3^. Based on its expression in the LPM, we believe that *Cer1* has replaced *Lefty* as the repressor of Nodal signaling in the left LPM of veiled chameleon embryos (Fig. 6).

One of the most interesting findings that sets veiled chameleons apart from avians is the apparent mode for the initial break in L-R asymmetry. We originally observed these morphological changes via live imaging, which revealed dynamic changes on the left side of the embryo. The break in molecular L-R symmetry appears to be triggered by a morphological change in the embryo at stage 10. The posterior neural tube tilts such that the midline becomes misaligned with the blastopore slit. We showed that these morphological changes are highlighted by *Shh* and *Chordin* midline expression. Interestingly, although not previously noted, similar molecular asymmetry involving different markers is also evident in turtle embryos (see Fig. 4 q of Kajikawa et al. 2020 and Fig. 3 G of Coolen et al. 2008)^3,39^. The morphological changes evident in veiled chameleons appear to be fundamentally different from avians and even-toed ungulates since there is no primitive streak and no node in this squamate reptile. In veiled chameleons, *Shh* and *Chordin* are expressed symmetrically within the floor plate and notochord, and provide a readout of morphological asymmetry, rather than drive the process. Furthermore, these morphological changes occur before any of the Nodal cascade markers are asymmetrically expressed (Fig. 6).

We hypothesized that although neither *Nodal1* nor *Nodal 2* is expressed at these stages, their proteins may be present and active. pSmad2 provides a read-out of active Nodal signaling, and, similar to *Nodal* gene expression, we could not detect pSmad2 until later stages, when it was limited to the left LPM. Thus, Nodal proteins are not the drivers of morphological asymmetries in veiled chameleon. Subsequently, asymmetric pSmad2 expression persists slightly longer than either of the *Nodal* transcripts, suggesting the retention of active Nodal protein at later stages in embryogenesis, after the transcript is downregulated.

Overall, it still remains to be determined what triggers these morphological changes and the initiation in the break in L-R symmetry in veiled chameleon. Several avenues for future study include exploration of the Marangoni flow^40^ and its effects on collective cell movements, calcium signaling, establishment of planar cell polarity, as well as cellular chirality changes. Calcium has been shown to play a critical role in establishing L-R patterning in vertebrate embryos that utilize motile cilia, but its role is poorly understood in embryos that establish L-R patterning without the use of motile cilia in the LRO, like chicken embryos ^41–43^. Planar cell polarity, as well as chiral changes in cell shape are known to play important roles in L-R patterning in multiple organisms but remain unexplored in reptiles^44^. The fact that veiled chameleon embryos are pre-gastrula at oviposition, makes them an ideal model for exploring the earliest morphological and molecular mechanism underpinning the breaking of symmetry and evolution of L-R patterning in squamates.

## METHODS

### Animal husbandry

All animal experiments were conducted in accordance with the Stowers Institute for Medical Research Institutional Animal Care and Use Committee approved protocol 2020-115. Veiled chameleon husbandry was performed in our Reptiles and Aquatics Facility as described previously^7–9^ following the protocols which are publicly available here: dx.doi.org/10.17504/protocols.io.bzhsp36e.

### Data availability

All original data underlying this manuscript is available and can be accessed from the Stowers Original Data Repository at http://www.stowers.org/research/publications/LIBPB-XXXX.

### Embryo collection and fixation

Veiled chameleon eggs were collected at oviposition in the Reptiles and Aquatics Facility at Stowers Institute for Medical Research. Eggs were incubated in deli cups with moist vermiculite at a constant temperature of 28 °C. Late-stage embryos were harvested in 1X Phosphate Buffered Saline (PBS), decapitated, and fixed overnight in 4% Paraformaldehyde (PFA) in 1X PBS at 4°C. After washing in 1X PBS, organs were isolated and imaged. The lungs were isolated from fresh embryos intact and perfused with 4% PFA through the trachea, which inflated them. The trachea was tied off with a hair, and the entire lung was fixed overnight in 4% PFA at 4°C prior to imaging.

To collect early-stage embryos, eggs were incubated for 50-70 days post oviposition (dpo). The eggs were cleared of large particles and wiped with RNaseZap wipes (Invitrogen AM9786) to minimize RNAse contamination. Clean eggs were candled to determine the position of the embryo under the leathery shell. We used fine scissors to cut a segment of the shell around the embryo and separate the embryo (attached to the shell) from the rest of the egg. The embryos were further separated from the shell and dissected out of the membranes in room temperature (RT) Tyrode’s solution, made in DEPC-treated water. Subsequently the embryos were fixed overnight at 4 °C in 4% PFA in DEPC 1X PBS, then dehydrated through an ascending methanol series into 100% methanol, and stored at −20 °C for future analysis, including RNA *in situ* hybridization, immunofluorescent antibody staining and immunohistochemistry.

Stages of embryonic development were determined and reported in this paper after Dufaure and Hubert (1961)^45^ and Diaz et al. (2019)^10^, with slight adaptations (Supplementary Dataset 1).

To obtain kidney cells, a 95dpo embryo was dissected in sterile conditions. Both kidneys were extracted, minced with a scalpel, treated with 25% trypsin for 5 minutes, and then further dissociated into near-single cell suspension. The cells were plated into a 35mm MatTek glass bottom culture dish (P35G-1.5-14-C) in the following culture media: DMEM/F-12 (Gibco 10565-018), 10% fetal bovine serum, 1X antibiotic antimycotic solution (Sigma Millipore A5955-100ML) and 1μg/ml gentamycin solution (VWR 97062-974) and allowed to attach overnight. The following morning the media was refreshed.

### Immunohistochemistry and immunofluorescent staining

Embryos were bleached in Dent’s bleach for 2 hours at room temperature after which they were rehydrated through a descending methanol series into PBS. Embryos were then thoroughly rinsed in PBS, incubated in blocking solution (PBS, 1% goat serum, 0.1% Tween-20) for 1hour at RT, followed by incubation with antibodies [MF-20, 1:25 (Developmental Studies Hybridoma Bank); pSMAD2, 1:500 (Cell Signaling Technology 3108S); and Arl13b 1:2000 (Proteintech 17711-1-AP)] in blocking solution overnight at 4°C. After washing thoroughly in blocking solution (3 x 20 minutes), the embryos were incubated in species-specific secondary antibodies for 2 hours at RT. For immunofluorescence, secondary antibodies were conjugated to fluorophores, and DAPI was added to detect nuclei. We used an HRP-conjugated secondary antibody to detect for MF-20 staining. To remove excess antibody, embryos were washed 3 x 20 minutes in 1X PBS with 0.1% Triton-X. Embryos stained with pSAMD2, were washed overnight to decrease background staining. Fluorescent embryos were equilibrated through an ascending glycerol seies and mounted in VECTASHIELD mounting medium (Vector Laboratories H-1200-10) between two coverslips to facilitate imaging of both the ventral and dorsal sides. MF-20 antibody labeling was developed using the DAB substrate kit for peroxidase staining (Vector Labs SK-4100).

Kidney cells were plated on 35mm MatTek glass bottom culture dishes (P35G-1.5-14-C), and then fixed after live imaging, in 2% PFA for 20 minutes, and the antibody staining, and subsequent imaging proceeded in the dish.

### Imaging and image processing

Brightfield images of wholemount embryos were taken on a Leica MZ9.5 dissecting microscope, with a Leica DFC550 camera. Manual Z-stacks were taken of each embryo, and in-focus projections were generated using Helicon Focus 5.3 software. For larger embryos and organs, manual Z-stacks were coupled with manual tiling. In-focus tiles were then stitched using Photoshop.

Brightfield images of sectioned embryos post RNA *in situ* hybridization were acquired using a Lumenera Infinity3-3URC color CCD (2.8MP) camera, running Micro-Manager ver. 1.4^46^ acquisition software on a Zeiss Axioplan 2ie upright microscope with HAL100 (100W halogen) illumination.

Brightfield images of sectioned embryos stained with Hematoxylin and Eosin (H&E) were acquired with a Hamamatsu Flash 4.0 sCMOS camera on an Olympus VS120 Whole Slide Scanner with 100 Slide Loader. Samples were illuminated by an UCB Transmission Lamp. The system was equipped with a neutral density filter for transmitted light. H&E samples were imaged with an UPLSAPO 20x objective, NA 0.75 and illuminated with an UCB Transmission Lamp.

Fluorescent images of pSMAD2 stained embryos were obtained on a Zeiss LSM-700 laser scanning confocal microscope including a fully motorized upright Axio Imager.Z2 base, FixGate main beam splitter, automatic pinhole, continuously variable secondary dichroic beam splitter (VSD) to separate emission between 2 photomultipliers (PMT) with individual emission filters. Images were acquired with 405nm (5mW) and 488nm (10mW) excitation lasers with short pass 490 nm, band pass 490-555 nm emission filters and a Zeiss Fluar 5x objective lens, N.A. 0.25. Gain was set to 588 for 405, 720 for 488.

Fluorescent images of Arl13b stained cells were acquired with a Zeiss LSM-700 laser scanning confocal microscope including a fully motorized upright Axio Imager.Z2 base, FixGate main beam splitter, automatic pinhole, continuously variable secondary dichroic beam splitter (VSD) to separate emission between two photomultipliers (PMT) with individual emission filters. Images were acquired with 405nm (5mW), 488nm (10mW), and 555nm (10mW) excitation lasers with short pass 490 nm, band pass 490-555 nm and band pass 505-600nm emission filters and a Zeiss Plan-Apochromat 20x objective lens, N.A. 0.8. Gain was set to 596 for 405, 559 for 488 and 544 for 555.

ImageJ was used to create maximum projections of imaged embryos and cells, and Photoshop and Illustrator were used to adjust brightness of fluorescent images, and further process them for the manuscript figures. When images were rotated for a display in anterior-posterior direction, missing background color at the edges of the images was filled in with Photoshop.

### Image normalization and fluorescence intensity measurement

pSMAD2 immunofluorescence images were normalized to DAPI as follows. In NIH ImageJ^47^, Z-stack images were sum projected, background subtracted, and subjected to a Gaussian blur of radius 2. From these images, a pixel-by-pixel ratio image of immunofluorescence to DAPI was calculated. The background region outside the embryo was cropped from the ratio image using a mask generated by performing an intensity threshold on the DAPI channel, with “fill holes” enabled. To quantify relative pSMAD2 staining across the embryo, normalized intensity profiles were generated from ratio images from line regions of interest (ROI) of width 200 pixels. A custom image analysis macro and instructions are available at Stowers Original Data Repository.

### RNA collection and cDNA synthesis

33 embryos from a single clutch, ranging from stage 7 of development to 8 somites were collected as described above. Embryos were pooled, and frozen at −80°C with minimal liquid, until further processing. RNA was extracted using QIAGEN RNeasy Plus Mini Kit (74134), and purified RNA was submitted to Cold Spring Harbor for sequencing.

We used SuperScript III First-Strand Synthesis System for RT-PCR (Invitrogen 18080-051) for cDNA synthesis, utilizing 100ng of RNA per reaction. We caried out both Oligo(dT) and randomer hexamers reactions. The two reactions were mixed, and that cocktail was used to clone gene fragments for RNA *in situ* hybridization probes (Supplemental Dataset 1).

### Transcriptome

RNA libraries were sequenced using the PacBio Sequel-II system at Cold Spring Harbor. Raw subread bams were then used for consensus (CCS) generation, primer removal, demultiplexing, refining, and clustering using PacBio’s established IsoSeq v3 pipeline (version 3.2.2). These clustered reads were then polished to generate a fasta file containing only high-quality isoforms with predicted accuracies >= 0.99.

To generate a final set of non-redundant transcripts for gene discovery and annotation, the IsoSeq high-quality isoforms were run through Cogent (version v6.0.0) following the recommended protocol when no reference genome assembly is available. Cogent was used to partition input sequences into gene families based on k-mer similarity (using a k-mer size of 30) and reconstruct the coding region sequence for each gene family. The Iso-Seq high-quality isoform sequences were then aligned with minimap2 to a reference made from the reconstructed coding region sequences. From this alignment, cDNA Cupcake (version 10.0.1) was used to filter away isoforms with low counts of aligned full-length reads and isoforms with degraded 5’ exons. Gene annotations were then assigned to each gene family coding region sequence using blastn megablast (BLAST 2.10.1+) with an e-value cutoff of 0.1 against the blast nt database and taking the single best blast hit.

The transcriptome shotgun assembly data was submitted to the NIH Sequence Read Archive (SRA: SRR22257482) and deposited at DDBJ/EMBL/GenBank and is available under the accession code GKDP00000000.

### Molecular phylogenetic analysis

We used known orthologous sequences of *Nodal1* and *Nodal2* genes to identify corresponding transcripts in the veiled chameleon transcriptome. We used open reading frames present in the transcripts as the presumed peptide sequences for veiled chameleon Nodal1/2 proteins. The species and exact protein sequences selected for phylogenetic analysis were from Kajikawa et al., 2020 ^3^. Protein sequences were retrieved from https://www.uniprot.org/^48^ and https://www.ncbi.nlm.nih.gov/ ^49^. We used the https://ngphylogeny.fr/ web interface for all steps required to generate the phylogenetic tree ^50^. The multiple alignment was performed using MAFT with “linsi” option selected ^51^. TrimAI was then used for alignment curation, with 0 gap threshold ^52^. We used FastTree for tree inference with 1,000 bootstrap replicates ^53–55^. The resultant tree was displayed, managed and annotated using The Interactive Tree of Life web interface (https://itol.embl.de/)^56^. Branch lengths (mutations per cite) are represented by numerical values.

### RNA in situ hybridization probes

PacBio mRNA sequencing results, as well as published transcriptome results^15^ were used to design probes of 350-800 bp in length (Supplementary Dataset 2). Fragments were amplified from cDNA and cloned into pCRII-TOPO (*Pitx2, FoxJ1, Dnah11, Lefty*), pCR-Blunt II-TOPO (*Nodal2, Cer1, Chordin*) or pGEM-T Easy (*Nodal1*). *Shh* and *Fgf8 in situ* plasmids were previously published^57^.

### RNA in situ hybridization

RNA in situ hybridization was performed using the following protocol. Day1: Embryos were rehydrated through a descending methanol series from 100% methanol to PBT (PBS + 0.1% Tween-20), followed by further washes in PBT (2 X 5 min). The embryos were then incubated in 10μg/ml Proteinase K in PBT for 7 minutes at RT, without rocking, followed by refixation in 4% PFA + 0.1% glutaraldehyde for 20 min. After washing in PBT (2 X 5 min), the embryos were incubated in 50/50 PBT/Prehybridization mix (50% formamide, 5XSSC pH 4.5, 2% SDS, 2% Blocking Reagent, 250 μg/ml torula RNA, 100 μg/ml heparin) for 5-10 min, followed by prewarmed (68-70 °C) prehybridization mix for 10 minutes at 68-70 °C, then 1 hour prehybridization mix at 68-70 °C and finally prehybridization mix plus digoxygenin-labeled riboprobes (~1 μg/ml) overnight at 68-70 °C.

The next day, embryos were rinsed in Solution X (50% formamide, 2X SSC pH 4.5, 1% SDS) at 68-70 °C, followed by 4 X 30 minutes washes in Solution X at 68-70 °C, then prewarmed solution X/MABT (1X MAB – Maleic acid, 0.1% Tween-20) 50/50 for 5-10 min. Embryos were then moved to RT, rinsed 3 X 5 minutes in MABT, followed by 2 X 30 minutes washes in MABT. The embryos were then incubated in 2% Blocking reagent (BR) in MABT for 1hour, followed by 20% Lamb Serum, 2% BR/MABT for 1-2 hours, and then overnight in 20% Lamb Serum, 2% BR/MABT plus anti DIG-AP antibody (1:2000).

The next day, embryos were washed 3 X 5 minutes MABT, followed by 7 X 45min MABT, 4 X 10 minutes in NTMT (100mM NaCl, 100mM Tris pH 9.5, 5mM MgCl2, 1% Tween-20, 2mM levamisole) and then finally NTMT + BCIP/NBT – at RT 1-2 hours before continuing overnight in NTMT + BCIP/NBT at 4 °C.

Color development continued at RT until the desired darkness of the substrate had been obtained. The total development time is typically 24-72 hours, depending on the probe and NTMT + BCIP/NBT solution needs to be changed regularly. The color reaction was stopped with incubation in PBS and then embryos were stored long-term in 4% PFA at 4 °C.

### Histological stains

After the whole mount embryos were imaged post RNA *in situ* hybridization, they were processed in 30% sucrose in 1X PBS for 1-2 hours at RT, or until embryos sunk to the bottom of the tube. Then the embryos were equilibrated into Tissue-Tek O.C.T. Compound (VWR #25608-930) and embedded for frozen sectioning. Embryos were sectioned transversely to be 10 μm on a Leica or NX70 Cryostar cryostat, postfixed, stained with nuclear fast red for improved visualization of tissue morphology, dehydrated in ethanol and xylene series, and cover slipped for imaging and long-term storage.

To best determine morphological features in embryos, embryos were dissected as described, fixed overnight and dehydrated in ascending ethanol series into 70% ethanol. Embryos were embedded in paraffin and sectioned transversely to be 5 μm. Embryos were subsequently stained with H&E using standard techniques, dehydrated in ethanol and xylene series, and cover slipped for imaging and long-term storage.

### SEM and TEM imaging

For SEM imaging embryos were fixed in 4% PFA in 1X PBS and processed for SEM as previously described in Jongebloed et al., 1999 ^58^ using tannic acid, osmium tetroxide, thiocarbohydrazide and osmium tetroxide (TOTO) to enhance conductivity before dehydration through a graded series of ethanol, followed by drying in a Tousimis Samdri 795 critical point dryer. Dried samples were then mounted on stubs, coated with 4 nm gold palladium in a Leica ACE600 coater and imaged in a Zeiss Merlin SEM at 20 kV with SE2 detector.

For TEM analysis, embryos were prefixed with 2.5% paraformaldehyde and 2% glutaraldehyde in 50 mM sodium cacodylate containing 1% sucrose (pH 7.4). The samples were post fixed in 2% OsO4. After dehydration through a graded ethanol series, samples were infiltrated and embedded in Epon resin (EMS, Fort Washington, PA). Ultrathin (60-80nm nm) sections were cut with a diamond knife and collected on single-slot copper grids. Grids were post-stained with 2% uranyl acetate and 1% lead citrate. Images were acquired on a FEI transmission electron microscope (Tecnai Bio-TWIN12, FEI) at 80kV.

### Live Imaging

Kidney cells, cultured in 35mm MatTek glass bottom culture dish (P35G-1.5-14-C) as described above, were imaged with a Nikon Apochromat TIRF 60x objective with N.A. 1.49 with an Orca Flash 4.0 sCMOS at 100fps at full resolution on a Nikon Eclipse Ti microscope equipped with a Yokagawa CSU W1 10,000 rpm Spinning Disk head with 50um pinholes. Samples were illuminated with transmitted white light LED light source. Fluorescent 2 μm beads (Polysciences Fluoresbrite^®^ YG Microspheres 2.00μm 18338-5) were added for better movement visualization.

Live embryos were dissected as described above and transferred to culture media containing DMEM/F-12 (Gibco 10565-018), 10% fetal bovine serum, 15% chick embryo extract^59^, 1X antibiotic antimycotic solution (Sigma Millipore A5955-100ML) and 1μg/ml gentamycin solution (VWR 97062-974)). Embryos were incubated in media and 5-SiR-Hoechst STED DNA dye^60^ (1:100 dilution) for 2 hours at 28 °C, and then mounted in the same media into an uncoated 35 mm glass bottom MatTek dish No. 1.5 (P35G-1.5-10-C) for imaging. Timelapse data were acquired on a Zeiss LSM-780 laser scanning confocal microscope equipped with a PMT, using a Plan-Apochromat 10x objective lens (N.A. 0.45). Z stacks were collected with 10μm spacing. Images were taken every 20 minutes at 30 °C to speed up development.

### Video processing

Timelapse data were processed in ImageJ as follows. Movies were corrected for intensity changes over time (due to increasing dye uptake) using the histogram matching method^61^. xy drift was corrected by applying the rigid body method for image registration to a max projected version of the timelapse and applying the resulting displacement and angle corrections to the full data set (using the stackregJ plugin available at Github.com/jayunruh/jayplugins and based upon Thevenaz et al., 1998^62^). The resulting timelapse data were then rotated in ImageJ using the standard “Rotate” command, such that the anterior-posterior axis was parallel to the vertical y-axis of the image.

Stacks were imported into Imaris software (Oxford Instruments) to generate final movies. Anterior views were generated using the “MIP” mode of Volume rendering and the “camera type” setting on “Orthogonal”, while cross sectional views were generated at indicated regions using the “Ortho Slicer” tool.

## ACKNOWLEDGEMENTS

The authors thank members of the Trainor lab for their insights and discussions throughout the course of this project. We are indebted to Rick Kupronis, David Jewell, Alex Muensch, Nikki Inlow, Diana Baumann and Elizabeth Evans and the Reptile Facility for their care, husbandry and maintenance of our veiled chameleon colony. We would like to thank the support of the Stowers technology centers, particularly microscopy for the live imaging help, and Hannah Wilson and Nancy Thomas from histology for paraffin sections and histological stains, as well as the bioinformatics team for assembling a de-novo transcriptome. This work was supported by an Emerging Model Organism grant from the Society for Developmental Biology (N.A.S) and the Stowers Institute for Medical Research (P.A.T).

