## Supplementary Materials for "Morphological changes and two *Nodal* paralogs drive left-right asymmetry in the squamate veiled chameleon (*C. calyptratus*)"

**Supplementary Dataset 1.** Embryo staging, based on (Dufaure and Hubert, 1961), adjusted for veiled chameleon-specific developmental features.

| Stage (Dufaure and Hubert, 1961) | Description |
|---|---|
| *Gastrulation* | |
| 7 | The embryo is mostly circular. The blastopore forms a canal, with the posterior end elevated over the plane of the embryo on the ventral side. The ventral opening is to the anterior, appears concave and relatively narrow. The dorsal opening of the blastopore is also narrow. The dorsal blastoporal lip is concave, with the opening towards the posterior, reminiscent of letter "C". |
| 7+ | The embryo elongates slightly and appears ovoid. The blastopore opening broadens, and the posterior blastoporal lip becomes reminiscent of letter "M". |
| *Neurulation* | |
| 8 | Prechordal plate appears and can be distinguished as a slight anterior invagination on the ventral side. Ventrally, the blastopore has a wide concave rectangular opening towards the anterior. On the dorsal side the blastopore is concave, with the opening towards the posterior. The overlap between the ventral and dorsal lips of the blastopore narrows. |
| 10 | The blastopore opening has migrated and is open dorso-ventrally. The axial mesoderm is distinct, with mesodermal groove beginning to form. Early evidence for the formation of the head process and the foregut in the anterior. |
| 11 | The blastopore opening is closing and becomes reminiscent of a slit. The mesodermal groove is distinct. The head process and the foregut in the anterior are becoming more prominent. |
| 16 | 2 somite pairs. The head process and the foregut in the anterior are becoming further distinct. On the dorsal side, neural folds are formed. The blastopore opening is closing, and the posterior tissue becomes elevated over the blastopore. |
| 17 | 3 somite pairs. The foregut pocket and the head prominence are fully formed |
| 18 | 4 somite pairs. The cephalic region is starting to tilt to embryo's right. |

DUFAURE, J. P. & HUBERT, J. 1961. Table de développement du lézard vivipare: *Lacerta (Zootoca) vivipara* Jacquin. *Archives Anatomie Microscopie Morphologie Experimental* 50, 309-328.

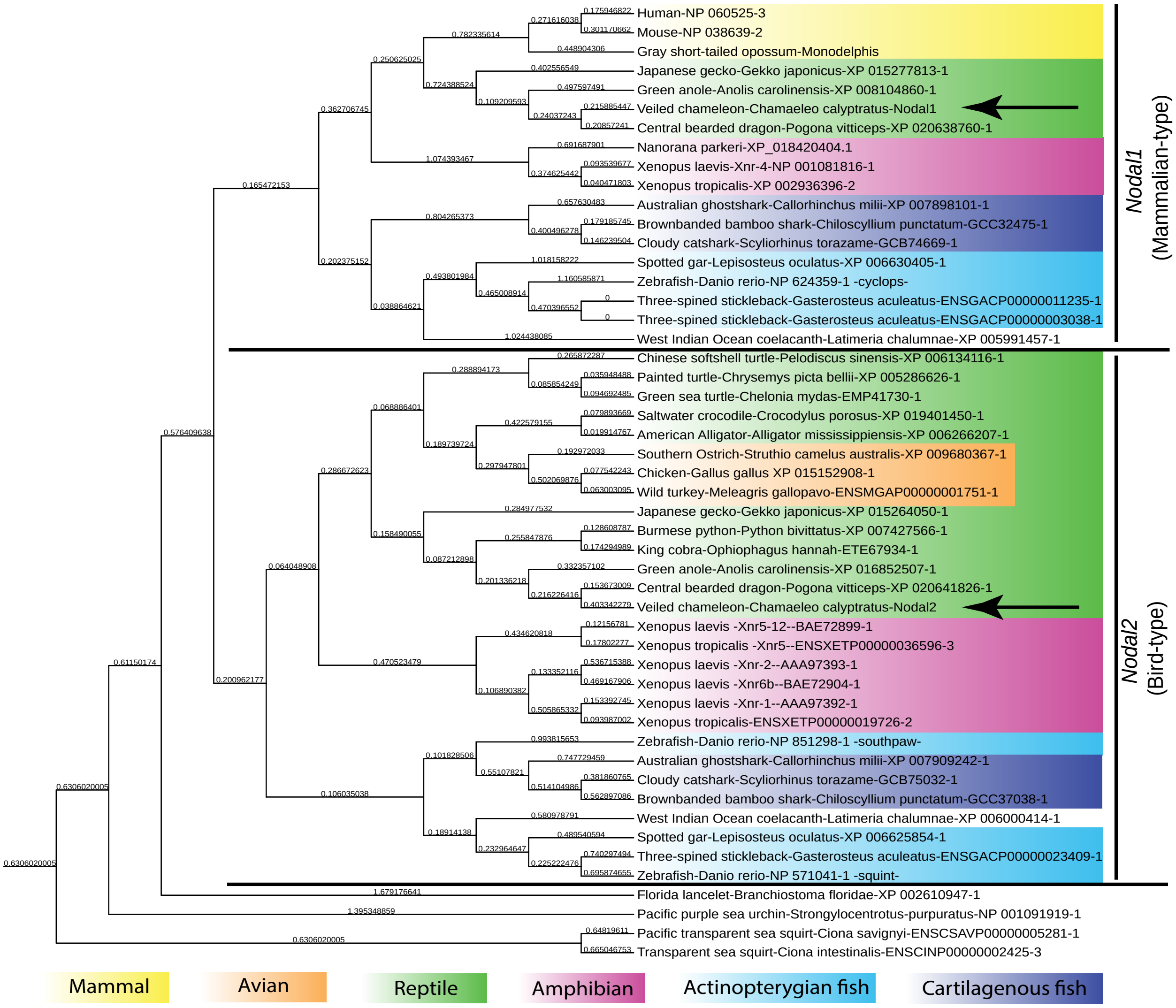


**Supplementary Figure 1. Molecular phylogeny of Nodal proteins.**

Veiled chameleon posseses two different transcripts, closely related to *Nodal* genes. Of the two ORFs, one sequence most closely aligns with Nodal1/mammalian type, and the other - with Nodal2/bird type (arrows). Background colors indicate taxonimic groups, in which selected species are categorized, and follow annotation laid out in Kijikawa et al., 2020. Not all phylogenetic branches are consistent with known phylogenetic relationships between species, likely skewed by mutation rate and and limited sequence input. The numerical values represent branch lengths (mutations per site).

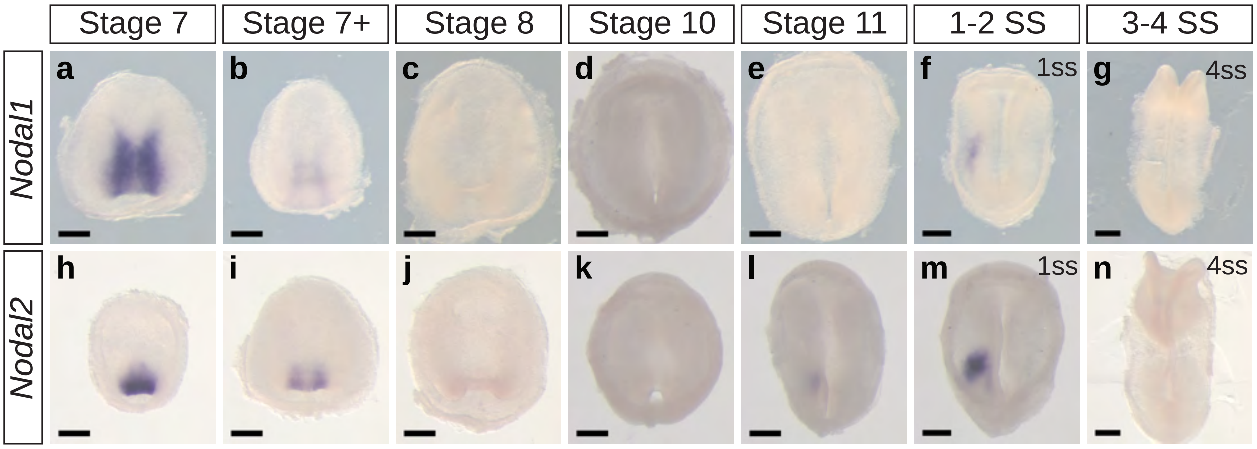


**Supplementary Fig. 2 | Dorsal view of *Nodal1* and *Nodal2* expression** All embryos are presented in dorsal view. Ventral view is available in Fig. 2. **a-g.** Whole mount RNA *in situ* hybridization for *Nodal1* expression. **h-n.** Whole mount RNA *in situ* hybridization for *Nodal2* expression. All scale bars are 200 µm.

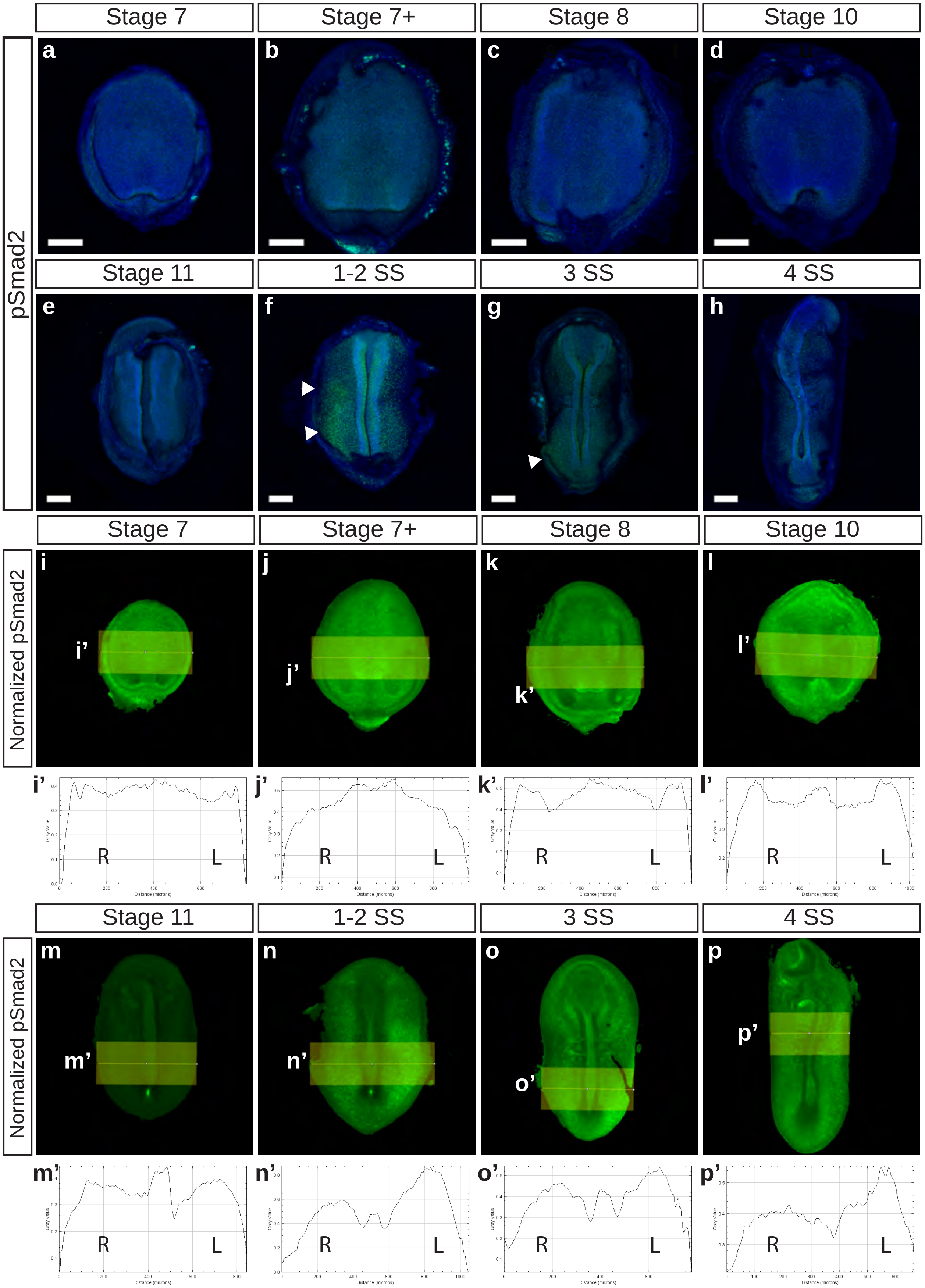

Stage 7
Stage 7+
Stage 8
Stage 10
pSmad2
a
b
c
d
Stage 11
1-2 SS
3 SS
4 SS
e
f
g
h
Normalized pSmad2
i
j
k
l
i'
j'
k'
l'
R
L
Gray Value
Distance (microns)
m
n
o
p
m'
n'
o'
p'

**Supplementary Fig. 3 | pSmad2 staining reveals Nodal1 and Nodal2 activity in veiled chameleon embryos.** Embryos in **a-h** are dorsal view, and embryos in **i-p** are ventral view of embryos in Fig. 2 **o-v. a-h.** The presence of active Nodal signaling was evaluated through antibody staining for pSmad2 (Green), with nuclear Dapi staining in blue. White arrowheads denote areas of pSmad2 enrichment visible in this view. **i-p.** pSmad2 staining normalized to Dapi staining. **i'-p'** intensity analysis between left and right sides on embryos, imaged in ventral view and normalized to Dapi. Areas analyzed as indicated in **i-p**. R (right) and L (left) denote the equivalent side of the embryo in the image analyzed. All scale bars are 200 µm.

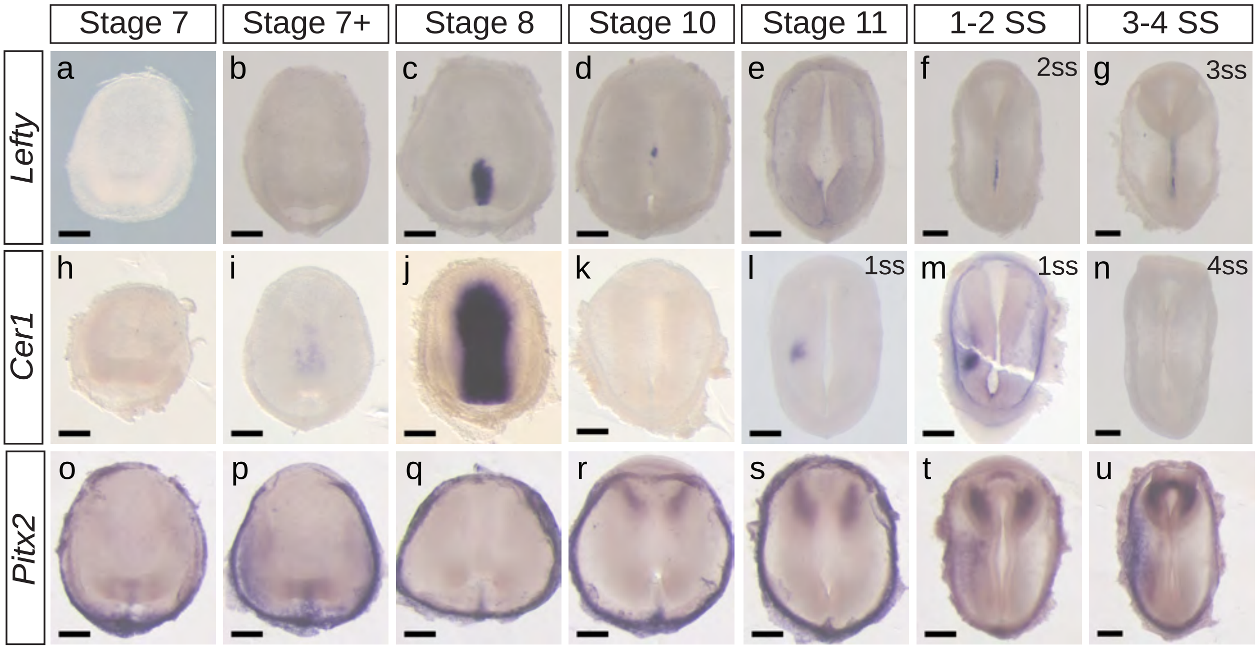


**Supplementary Fig. 4 | Dorsal view for expression patters of key members of the Nodal cascade in veiled chameleon embryos.** All embryos are presented in dorsal view. Ventral view is available in Fig. 3. **a-g.** Whole mount RNA *in situ* hybridization for *Lefty* expression. **h-n.** Whole mount RNA *in situ* hybridization for *Cer1* expression. **o-u.** Whole mount RNA *in situ* hybridization for *Pitx2* expression. All scale bars are 200 µm.

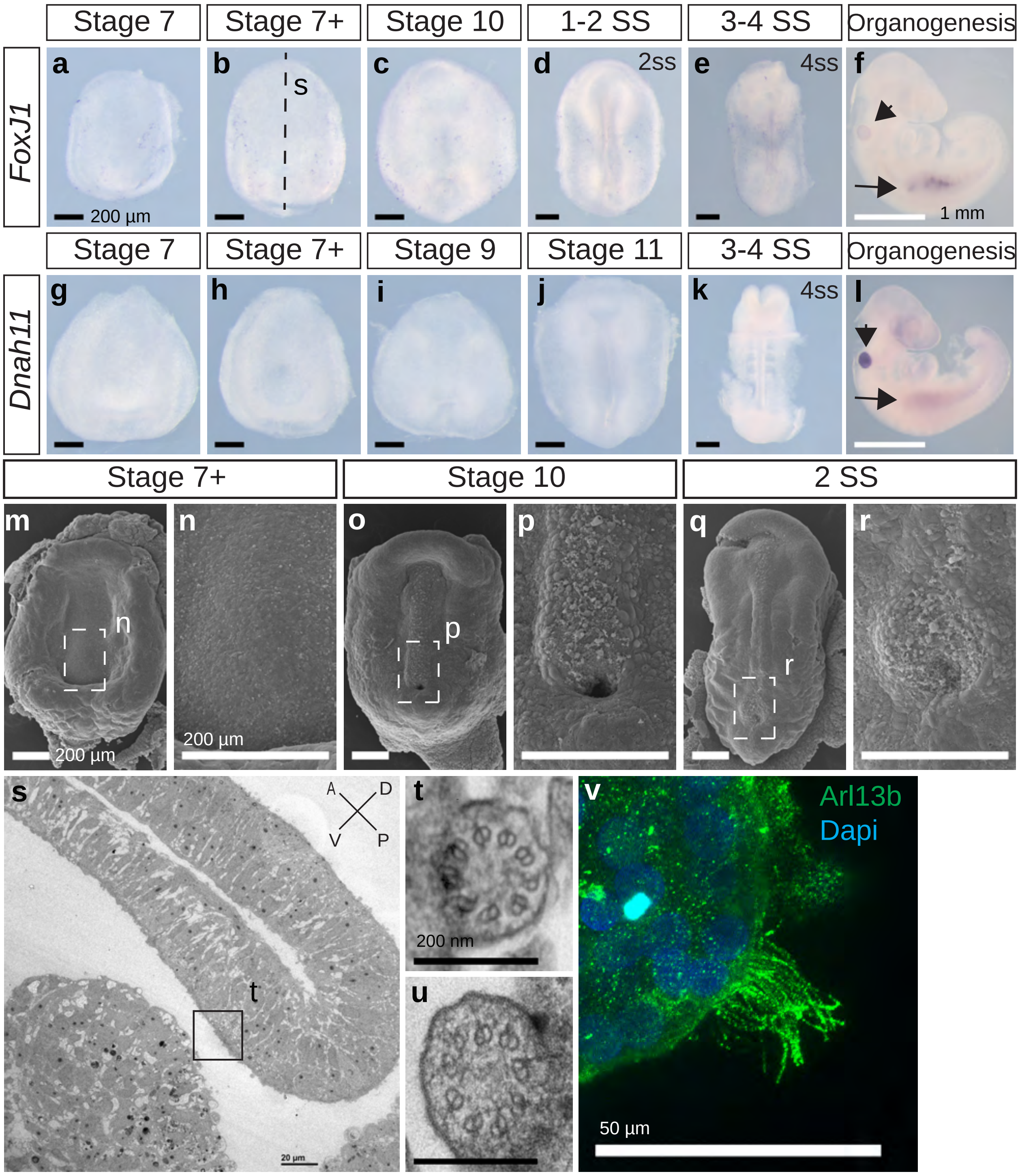


**Supplementary Fig. 5 | Veiled chameleon embryos lack motile cilia during L-R asymmetry establishment.** All embryos are presented in ventral view, unless otherwise noted. Dorsal view available in Fig. S6, where possible. **a-f.** Whole mount RNA *in situ* hybridization for *FoxJ1* expression. **g-l.** Whole mount RNA *in situ* hybridization for *Dnah11* expression. **f, l**. Arrowheads indicate gene expression in otic vesicle, arrows indicate gene expression in the kidney – both areas know to have motile cilia. Black scale bars in **a-e** and **g-k** are 200 µm. White scale bars in **f, l** are 1mm. **m-r.** SEM visualization of embryos. Dashed rectangular areas in **m, o, q** are enlarged in **n, p, r.** All scale bars in **m-r** are 200 µm. **s-u.** TEM images. **s.** Sagittal section through an embryo, as indicated with a dashed line in **b** for a stage-matched embryo. Embryo orientation is as indicated, with blastoporal lip located dorsally. Scale bar 20 µm. Boxed region indicates exact location of the cilium, depicted in **t. t, u.** Cross sections of two cilia, located near the blastopore, as indicated in **s.** Both cilia have 9+0 structure and lack axonemal dyneins, pointing to their nature as primary, non-motile cilia. Scale bars are 200 nm. **v.** Fluorescent immunostaining of motile multicilia in dissociated kidney cells. Ciliary marker Arl13b is in green, nuclei labeled with Dapi in blue. Video of ciliary movement is available in Supplementary video 1. Scale bar is 50 µm.

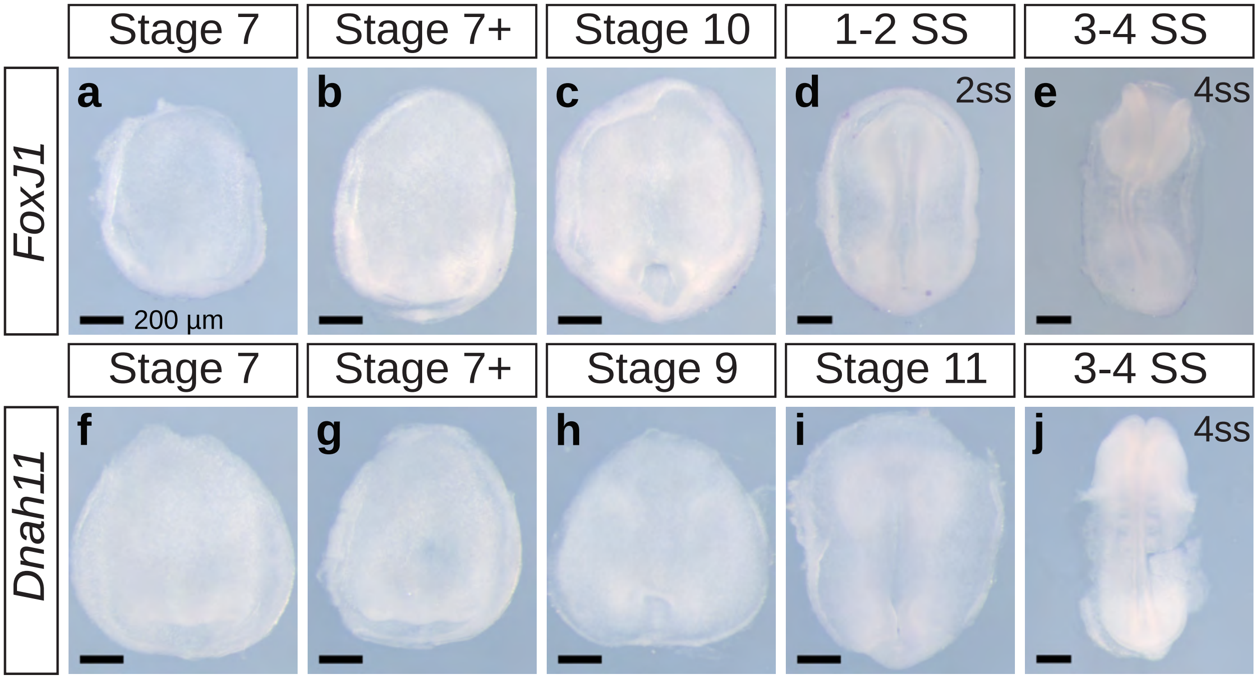


**Supplementary Fig. 6 | Dorsal view of *FoxJ1* and *Dnah11* ciliary markers expression.** All embryos are presented in dorsal view. Ventral view is available in Fig. S5. **a-e.** Whole mount RNA *in situ* hybridization for *FoxJ1* expression. **f-j.** Whole mount RNA *in situ* hybridization for *Dnah11* expression. Scale bars are 200 µm.

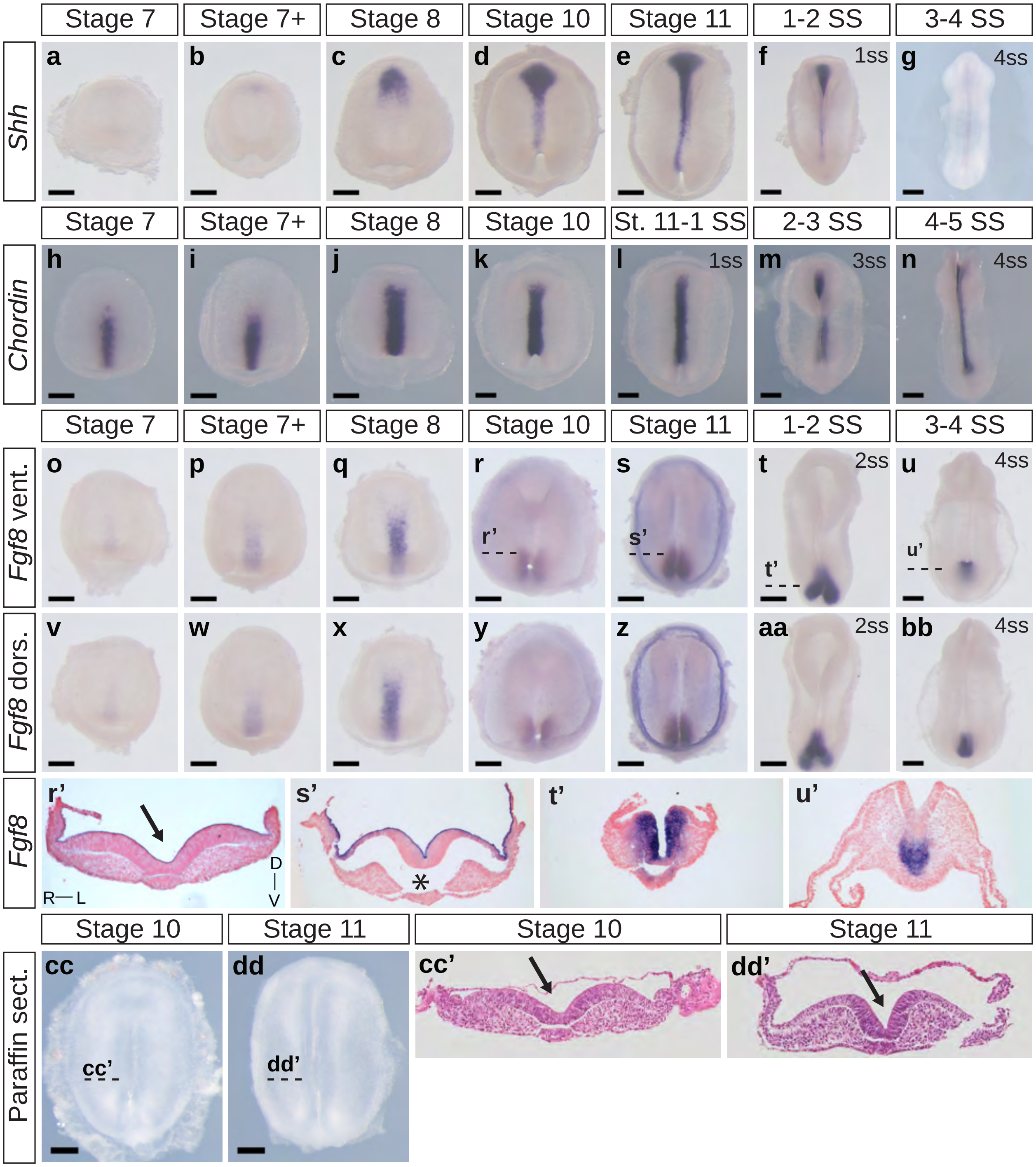


**Supplementary Fig. 7 | Morphological changes in the embryo visualized through H&E staining, *Shh, Chordin* and *Fgf8* expression. a-n.** Dorsal view of embryos from Fig. 4. **a-g.** Whole mount RNA *in situ* hybridization for *Shh* expression. **h-j.** Whole mount RNA *in situ* hybridization for *Chordin* expression. **o-u.** Ventral view of whole mount RNA *in situ* hybridization for *Fgf8* expression. **v-bb.** Dorsal view of embryos from **o-u. r'-u'**. Transverse sections of embryos from **o-u,** as indicated with dashed lines. Sections were counterstained with nuclear fast red for better visualization. **cc-dd**. Ventral view of embryos, sectioned transversely and stained with H&E. Approximate location of sections indicated with dashed lines. **cc'-dd'**. Transverse paraffin sections of embryos in **cc** and **dd**, stained with H&E. Section orientation is as indicated in **r'**. Arrows point to the medial hinge point of the neural tube. Asterisk reveals asymmetry in notochordal plate. All scale bars are 200 µm.

**Supplementary Dataset 2.** Veiled chameleon RNA *in situ* hybridization probe sequences

>*Nodal1*
TCTGGCTGAGGTTCGACTTCACCTGCCCAGTTCTGGAGAGAAATCTACCTTTAACAG
TGCCCCAATGTGGGTAGACCTGTATCACCAGCAGGAAATCCATTGCTCAAGTGGTCA
CAACTGCCCACACATTGTTCACATAGGCTCTTTTGAAGCGCCTCCATCATTGCCTTCA
AGCTGGATAGTTCTGGAAGTCACGGAACAGCTCTCAAAGTGGGTAGTGAACAGCAG
CTTAGTGGAGGAATCTATTCAGAAAACTCTAAGTGAGGAACACCAGCATAGCAAAC
ATTTGCCAAAGCAAGCAACAAGTTACTGTGGATCTATAGATAGAAAGGCATTCTTAG
TGCTTTTCTCTAGACTTTCCAAAGAAGAGAAAGAAAGAAATAGCTCAACTTTGCTCC
AGATAGTGAAAGACTCAAAATATCTTACACTGAAGAATCCAAAAGATGTGGTACCT
GTTTCGGGAACAAAGAGGTATCGGCGCCACAAGAAACTCCAAGGCTGGTCTTTAAT
GAGTTTGAAAGAACTCAAGGTCGAAAACCTATGCCATAGAGTGGACTTCTATGTTGA
TTTTGAGGAGATAGGCTGGGGCTCCTGGATTATATACCCCAAGAGATATAATGCCTA
CCGCTGCGAGGGCATGTGTCCTA

>*Nodal2*
ACCTCGTCCTTCTCGTAGTAAAGCATGGAGAGTGGGCTCAATTTGACTGGAGCGCAG
GCTAGGCAAGGCACCCGGTTGGGTTGGTAGAACTTCAGCAAGCTCTGCATATATGCA
TGGTTGGTTGGCTTAAAGGTTTCATCCACAGGTGTTGGGCACTCCCCCTCACAACGG
TATGCATTATACTTCTTGGGATGTAGGATCCACATTCCAAAACCAGTCTTCTCAAAG
TCCACAATCATATCAACTCTCTTGCACAGGGATTTATTTTCTTCAGCAAAGCCAAGC
ACGTCAGCGTCGCTCCCTTTGATCCTTTTCCTTTGCTTCCTGTTCCGTCGGTGCCTTCG
GCCCCTGGGCTCTTTGTGTGCGTCATCTGACATGACGTATTTGGACATCTCTACCACC
CTGATGAGACTTGATCTCTGATAGGGTTCCATCTGTTCTTTATCTTTGGAGAAGATAA
CCAGAAGGACTTGTTCTTTCCTGCTCTGCAACGGTACTTCTGGGGTTGGACTGAAGT
CACAGATGTTCTCAAACCCATTGGTGGGGCATCCACTGGTTCCTTCGTCATTGCCATT
ATCTAGGCAATTTGCAGTCCATTCCTGCTCCTGAACATCTAGGGTATCGTGACCTGA
G

>*Lefty*
GTTTCCCTTCTTGACAAGGTACATCATGGAAGTGGGGAACTTTCCACCACAGCACA
GTTTCTCTCTCCATAGCCAAAATGCCGCAACCAGTTCTTGGGTTGCCGGCAGCAACC
CATACAGTGGTAAGCCTGATAGCCAGCTGGCTCAATAATCCAATATTGAGTCCAAGT
TAACTCTCGGAAATTGATGTAATGTTCTTGCCGGCAGCAGGTGGTTTTGTCCGTTGAT
GCACCTTCTTTACAGTCTCCAGGTCCTCCATACTCTTCCAAGTTTATTGTATAAAGCA
CTAACTCAGGCTTGCCCAACGTCTTATCGGAGGGGTCCTGAGAAGTAAAGCGCACA
ACTCTGGCCATTTCTGAGGCATAGTTGCCTATTCTTTCTCCTTCAATCCAGACCTCCA
GAAGCATTGGCCCAGTTTTCTTCACTTTAAGCCAAAAATGTACAGCCTGGGTGACAT
CAAAATTCTTCCAGCCAGATTCCATAATTGGGACCAATCTGGAATCAATCAAAGTAG
TCCGGTTGGTTCCATTATCTCCAAGCTCTACCCAGTAGATGCTCACCCGGGCATTCA
AGACGGGTCTTTGAGATTGCCTGCTGGGCAGGTTTTTTATATCCAGGGGCTTCTTGA
AAAGTTTCAGCTCAGCCAT

>*Cer1*
CTGCAAACATCCTGGTACATTTCATTGATCTTGATTGGAAGGACAATCTCTTCAGAC

CTTGATTTGGTTTTCAACATGAACAGATCCCAGAACTTCTTGGCATCTTTTCGGAAGA
CCAGGTTTGCCTTTCTCTTAGCGTGCTGATGCGGAGGATGGTTTGAGGGGGTTTCTG
GGTATGAAGGTGGATCTGTGGGAATCCAGCTTTCCAAATCCTGAGCCATATGGCCAC
CAACATGTGGGAAAATTGTCCTGGACACTTCAGGTTCATCTTGTGTCTTGCTCTCTGT
TTCTGCCAAGACAGCTGCCACAAACGGATCAGGCTGTGCCAGATACCTCACTAACA
AATCCTGAGGCAACTCCCGACTTTGTTCTGCTCTCTCAGGCCATGAAATCAGCAGCA
ACAAATACAAGATGCAGGCCTTG

>*Pitx2*
GGATCCGTCCAAGAAGAAGAGGCAGCGGCGGCAGCGCACGCACTTCACCAGCCAGC
AGCTCCAGGAGCTGGAAGCCACCTTCCAAAGGAACCGCTACCCGGACATGTCGACG
CGGGAGGAGATCGCCGTCTGGACCAACCTCACCGAGGCCCGAGTCAGGGTGTGGTT
CAAGAACCGCCGGGCCAAGTGGCGGAAGCGGGAACGGAACCAGCAGGCGGAGCTG
TGCAAGAACGGCTTCGGGCCGCAGTTCAACGGCCTGATGCAGCCTTACGACGACAT
GTACCCGGGCTACTCGTACAACAACTGGGCGGCCAAGGGCCTGACGTCGGCCTCGC
TCTCCACCAAGAGCTTCCCCTTCTTCAACTCCATGAACGTCAACCCGCTCTCGTCGCA
GAGCATGTTCTCGCCGCCCAACTCCATCTCGTCCATGAGCATGTCTTCCAGCATGGT
CCCCTCGGCCGTGGCCGGCGTCCCGGGCTCGGGCCTCAACAGCCTCAACAACTTGAA
CAACCTGAGCAACCCTTCGCTCAACTCGGCCGTGCCCACGCCCGCCTGCCCTTACGC
CCCGCCGACCCCTCCTTACGTTTACCGGGACACGTGTAACTCCAGCTTGGCCAGCCT
GAGACTCAAAGCCAAGCAGCACT

>*Shh* [1]
TCCGGCTTCGACTGGGTCTATTACGAGTCCAAGGCTCACATCCACTGCTCGGTCAAA
GCCGAGAACTCAGTGGCAGCCAAATCCGGTGGCTGTTTCCCCGGAACAGCCTGGGT
GAACCTGGAGCAAGGAGGAACCAAGCTGGTGAAGGACCTCAACCCTGGAGACCGG
GTTTTGGCAGCGGACACCCAGGGCCGCCTGCTCTTCAGTGAATTCCTCACCTTCCTG
GACCAAGAGGAGGCCCCGATCCACAAACTCTTCTATGTCATTGAGACCCAAAGGCC
CCGGACACGTCTCTTGCTGACGGCGGCCCACCTACTCTTTGTGGCCTCACCCCAGAA
CCAGTCTCAACCCCAGCCCATTTTTGCCAGCCGTGTACAGCCAGGACAACATATCTA
CGTGCTCAGCCAAGGAGGTCAGACACTGCTAGAAGCTGCAGTGCACCGGGTTTCCC
TGCAAGAGGAAGCTTTGGGGGCCTACGCCCCACTGACCGCCCATGA

>*Chordin*
CATACCCTATGGGAGAGATCCAAGGGAAGATTATCAAGCATCGGGCCCTCTTTGCA
GAAACATTCAGTGCCCTCCTGACATCTGCGGACCCAACCCACCTTGGCATGGGTGGC
ATTGCTATGCTTACCTTGAGTGACACAGAGAACAACCTACATTTTGTCCTAGTGACC
AGGGGGCTTTTGGAGCCAGCTGACAAAAAATCCTCCTCGCTCCCCTTGAGGGTACAA
ATCCTGCACCAGGACAGAGTTCTGAGGGAAATGCTTGCCAATGTTACCTTGCAGGCC
TCTGACTTCGCAGAAGTGCTGACTGGACTTGGAAGTGAAGAGATGCAGTGGCTGGC
CCACGGGGCATTGAGGATTACAGCAGAGGTGGAGGGGAAATTCAGGCGCCAGATTG
CTGGACAGATCACCCCAAGGAGAAGTTGTGACACCCTACAAAGTGTGCTTTGTGGA
GGAGATGCTTTGATACCGACCCAGACAG

*>Fgf8*[1]
TCTAGTGCGGACCTACCAGCTGTACGGTCGGACCAGTGGGAAGCACGTGCAAATTCT
GGACAACAAGAAAATCAATGCCTTGGCAGAGGATGGAGACGCTCACGCCAAGCTCA
TTGTGGAGACTGACACCTTTGGAAGCCGAGTAAGAATCAAGGGTGCTGAAACTGGC
TTCTATATCTGCATGAACAAGAAAGGAAAGCTGATTGGCAAGAGCAATGGCAAAGG
CAAGGACTGTGTCTTTACAGAGATTGTGTTGGAGAACAACCACACAGCACTGCAGA
ATGCCAAGTATGAAGGGTGGTACATGGCCTTCACCCGAAAGGGGCGCCCCCGCAAA
GGCTCCAAGACTCGTCAACATCAACGTGAGGTGCACTTCATGAAGCGCTTA

*>FoxJ1*
CATTAGGCACAACCTCTCCTTGAACAAGTGCTTCATCAAAGTCCCACGAGAGAAGG
ATGAACCAGGAAAAGGTGGCTTCTGGAAAATTGACCCTCAGTATGCTGACAGACTG
ATGAACGGGGCATTCAAGAAGCGCAGGATGCCTCCAGTGCAAATCCATCCGGCTTT
CAGTGGACGAATGCAACAGGATGCCTGCTCCAGTTCTTCTGCTCAGCAGGCTGCAAT
CTCTTGGAAAAACAATGGCATCCTAAAAATCAACATGGAGTCTCAGCAGCTACTCA
AAGAATTTGAAGAAGTCACTAGTAGTGATCAAAACTGGAATCCAGCAGTGGATGGG
AAAATGAGCCATAAACGTAAGCAGCCTTTGCCAAAACGGATGTACAAGACTGCCCG
CCTCTCCAGCTCTCCCATGCTGACACAGGAAGAACAAACAGAGCTTGGATCTCTGAA
AGGTGACTTTGATTGGGAAGCTATCTTTGACACCACTTTAAATGCTGATTTTTCCACC
TTTGAAGATTTGGAGATCACGCCTCCTATTAGCCCAATAACCAGGGATGTAGATTTG
ACTGTGCATGGAAGACATATCGATTGCCCACAGGAGTGGTGCCCCACTGGGCAGGA
TTACGTTCTAACAGAGTCCAACCAGAACAGCTTAGACTTTGATGAAACCTTCATTGC
TACTTCTTTCCTCCAGCATCC

*>Dnah11*
CTGCCAGTCTTTTTGAAGTGGTCAGTCCAGATTACAAACAACTGAAACAGTGTCGCA
AGGAAATAACATTGCTGAAGGGAATGTGGGACATCAATATTTATGCAACAAGTAAC
ATCAGTGATTGGATTAAAAGCCCTTGGAGGGAGATTAGTATGGAGCAGATGGATGC
AGAACTGAGAAGATTTGCAAAGGAGTTGTGGGCACTGGATAAAGAAGTTCGCTCCT
GGAATGTATACACAAATCTGGAACTAACAATTAAAAACTTGTTGACATCGTTGAAG
GTTGTTACAGAGTTGCAGAATCCAGCCATGAGGGACAGGCACTGGCATCAGCTGAT
GGATGCAATAGGTATTCAGTTTTCAATAAGTGAAGATACAACATTGGCAGATTTGTT
AGCGCTGAAGCTCCACAAGATGGAAGATGATGTCAGAAACATTGTTGACAAAGCAG
TAAAAGAACTTGGGATTGAAAAGATTCTCACAGAAATCAACCAAATATGGGCTACA
ATGGAGTTTTGTTATGAAGAGCATTACAGGACCAGTGTTCCTTTGTTGAAAACAGAT
GAGCACCTTTTTGAGACATTAGACGATAACCAAGTTCAGCTGCAAACAGTTCTGCAA
AGTAAATATGTTGAGTATTTCATTGAGCAAGTTTCAAACTGGCAAAAAAAGCTAAAT
ATTGCAGACTCTGTAATTTTTCTTTGGATGGAAGTTCAGCGCACATGGTCTCATCTTG
AAAGCATTTTCATTGG

1 Diaz, R. E., Jr. & Trainor, P. A. Hand/foot splitting and the 're-evolution' of mesopodial skeletal elements during the evolution and radiation of chameleons. *BMC Evol Biol* **15**, 184 (2015).
